# Small-molecule activators of the *Staphylococcus aureus* ClpC/ClpP AAA+ protease

**DOI:** 10.64898/2026.04.09.717423

**Authors:** Timo Jenne, Vselovod Viliuga, Ulrike Uhrig, Britta Jehle, Merlin Schwan, Jürgen Kopp, Dirk Flemming, Elisabeth Seebach, Irmgard Sinning, Bernd Bukau, Axel Mogk

**Affiliations:** Center for Molecular Biology of Heidelberg University (ZMBH), DKFZ-ZMBH Alliance, Im Neuenheimer Feld 345, 69120 Heidelberg, Germany; Dept. of Biochemistry and Biophysics at Stockholm University and Science for Life Laboratory, Stockholm, Sweden; The European Molecular Biology Laboratory, EMBL, Meyerhofstraße 1, 69120, Heidelberg, Germany; Heidelberg University Biochemistry Center (BZH), Im Neuenheimer Feld 328, 69120 Heidelberg, Germany; Department of Infectious Diseases, Medical Microbiology and Hygiene, Medical Faculty Heidelberg, Heidelberg University, Heidelberg, Germany; University Hospital Heidelberg, Heidelberg, Germany

**Keywords:** Protein degradation, AAA protein, Hsp100, protease, chaperone, bacteria, ClpP, protein quality control, SAR, MoA

## Abstract

The central AAA+ ClpC/ClpP protease in Gram-positive bacteria is crucial for virulence and stress resistance and has been recognized as drug target. Natural cyclic peptides deregulate the essential *Mycobacterium tuberculosis* ClpC1 and cause cell death. Similarly, overactivated mutants of the non-essential *Staphylococcus aureus* ClpC homologue cause uncontrolled proteolysis and severe toxicity *in vivo*. However, no chemical modulators of *S. aureus* ClpC have been described. Here, using a biochemical high-throughput screen we identify eight chemically distinct *bona fide* small molecules that robustly stimulate ClpC ATPase and proteolytic activity *in vitro*. Structural, computational, and mutational analyses define two ligandable regulatory sites within the ClpC N-terminal domain (NTD) as compound targets: a conserved hydrophobic groove and an allosteric pArg1 pocket, both engaged in substrate recognition. These findings establish *S. aureus* ClpC as chemically targetable and provide mechanistic insight into its regulatory architecture, enabling future development and optimization of chemical probes to deregulate AAA+ protease control.

## Introduction

Recently, there has been a significant increase in hospitalized patients with difficult-to-treat infections caused by drug-resistant bacteria, a trend most pronounced for the pathogen *Staphylococcus aureus* (GBD 2021 Antimicrobial Resistance Collaborators 2024). While strains such as totally drug-resistant *Mycobacterium tuberculosis* (TDR-TB) evolved (Daneshi et al. 2025) and methicillin-resistant *Staphylococcus aureus* (MRSA) strains emerge at high frequency (Subramanian et al. 2025), the number of new antibiotics developed over the last 50 years is very small (Lewis 2020). Traditional antibiotic targets such as protein biosynthesis and DNA replication have been largely exhausted (Akunne et al. 2025), demanding the identification of new antibacterial targets for antibiotic development.

One promising new drug target is the central ClpC/ClpP protease in Gram-positive bacteria, a functional equivalent of the eukaryotic 26S proteasome (Sauer and Baker 2011). The ClpC/ClpP proteolytic machinery consists of two major components, the AAA+ (ATPase associated with diverse cellular activities) unfoldase ClpC and the barrel-shaped serin peptidase ClpP. ClpC consists of two ATPase domains (AAA1, AAA2), an N-terminal domain (NTD), and a coiled-coil M-domain (MD). In its active state, ClpC assembles into a hexameric ring and threads substrates in an ATP-dependent manner through its central translocation channel directly into the associated proteolytic chamber of ClpP (Azinas et al. 2025). To prevent unspecific unfolding and degradation events, ClpC activity and substrate specificity are tightly regulated. In its basal state, ClpC forms an inactive resting state composed of two half-spirals (Carroni et al. 2017; Jenne et al. 2025). In this resting state, ClpC exhibits low ATPase activity and cannot associate with ClpP. This assembly state is stabilized by head-to-head-interacting MDs and by the binding of the NTD to MDs and the AAA1 domain. ClpC activation involves hexamer formation mediated by various adapter proteins that bind to the NTD (MdfA, McsB) or both MD and NTD (MecA) (Massoni et al. 2025; Wang et al. 2011; Kirstein et al. 2007). Alternatively, polypeptides harboring a phosphoarginine (pArg) degron tag created by the McsA/McsB arginine kinase are specifically recognized by two conserved binding pockets (pArg1/pArg2) at the NTD, also triggering ClpC activation (Fuhrmann et al. 2009; Jenne et al. 2025; Trentini et al. 2016). Importantly, MecA and McsB target themselves for degradation in the absence of substrate, forming a negative feedback loop that restricts ClpC activity to the presence of substrates (Schlothauer et al. 2003; Kirstein et al. 2007).

ClpC/ClpP is crucial for stress resistance, virulence, and biofilm formation in various pathogens such as *Staphylococcus aureus*, *Listeria monocytogenes,* and *Streptococcus pneumoniae* (Bhandari et al. 2018; Frees et al. 2004; Ibrahim et al. 2005; Luong et al. 2011; Rouquette et al. 1998; Zhang et al. 2025). In mycobacteria, the homologous ClpC1/ClpP1P2 protease is even essential for cell viability (Lunge et al. 2020; Ollinger et al. 2012). Multiple screening campaigns identified *M. tuberculosis* ClpC1 and also ClpP1P2 as a potential drug target (Choules et al. 2019; Gao et al. 2015; Gavrish et al. 2014; Kazmaier and Junk 2021; Schmitt et al. 2011; Famulla et al. 2016). Additionally, deregulating ClpP from *S. aureus*, *Escherichia coli, Neisseria sp.*, and *Bacillus subtilis* has been shown to represent a viable strategy for antibiotic development (Brötz-Oesterhelt et al. 2005; Leung et al. 2011; Michel, K.H. & Kastner, R.E. 1985; Binepal et al. 2020).

One class of these antibiotics targeting ClpP are acyldepsipeptides (ADEPs) (Brötz-Oesterhelt et al. 2005). ADEPs deregulate ClpP by mimicking AAA+ partner docking by binding to ClpP H-sites, triggering formation of the active proteolytic site and opening of the ClpP entry pores (B.-G. Lee et al. 2010). This enables ClpP to autonomously degrade loosely folded proteins, including nascent polypeptides, causing toxicity (Brötz-Oesterhelt et al. 2005; Kirstein et al. 2009). Furthermore, ADEPs disrupt AAA+ partner-peptidase cooperation, thereby stabilizing native substrates, causing lethality in *M. tuberculosis* (Brötz-Oesterhelt et al. 2005; Famulla et al. 2016; Kirstein et al. 2009). ADEPs have been shown to be highly effective and even eliminate hard-to-kill persister cells (Conlon et al. 2013).

Molecules that target ClpC1 encompass several highly toxic natural cyclic peptides, including Cyclomarin A (CymA), lassomycin, rufomycin, and ecumicin (Choules et al. 2019; Gao et al. 2015; Gavrish et al. 2014; Schmitt et al. 2011). All these molecules bind to a conserved hydrophobic groove of the ClpC1 NTD, which functions as substrate binding site (Jagdev et al. 2023; Vasudevan et al. 2013; Wolf et al. 2020, 2019). Notably, the effects of cyclic peptides on ClpC1/ClpP1P2 activity vary. Peptides typically increase ClpC1 ATPase activity, yet can have both stimulatory and inhibitory impacts on proteolytic activities (Choules et al. 2019; Gao et al. 2015; Gavrish et al. 2014; Maurer et al. 2019; Taylor et al. 2022). Similarly, cyclic peptides differ substantially in their impact on the *M. tuberculosis* proteome, suggesting diverse modes of ClpC1 deregulation (Barter et al. 2026). Newly developed CymA-based degraders (BacPROTACs) enable targeted degradation of specific proteins of interest or even ClpC1 itself, representing alternative strategies to disturb protein homeostasis via ClpC1/ClpP1P2 (Morreale et al. 2022; Hoi et al. 2023).

Although ClpC homologues share high sequence homology, the previously described cyclic peptides are highly specific to mycobacteria (Choules et al. 2019; Gao et al. 2015; Gavrish et al. 2014) and do not target e.g. *S. aureus* ClpC. Yet, transplanting the NTD of *M. tuberculosis* ClpC1 to *S. aureus* ClpC leads to its sensitization towards CymA, and the fusion construct creates toxicity by uncontrolled proteolysis when expressed in *E. coli* in the presence of CymA (Maurer et al. 2019). This suggests that an induced, toxic deregulation of *S. aureus* ClpC/ClpP is possible, although ClpC and ClpP are not essential (Frees et al. 2004). Accordingly, expression of deregulated *S. aureus* ClpC mutants, which exhibit autonomous, uncontrolled activities, is highly harmful to *E. coli* cells (Carroni et al. 2017; Jenne et al. 2025). We therefore sought to identify small molecules that trigger persistent *S. aureus* ClpC/ClpP activation to disrupt protein homeostasis and conducted a high-throughput screen. We present the first examples of small-molecule activation of the *Staphylococcus aureus* ClpC AAA+ unfoldase *in vitro*. We identify eight *bona fide* compounds targeting two chemically ligandable regulatory sites of the NTD —the hydrophobic groove and the pArg1 pocket. Additionally, we clarified the roles of compounds in allosteric activation of ClpC through co-crystallography, computational approaches, and biochemically by site-directed mutagenesis. The identified compounds impair *S. aureus* cell growth, however, in a ClpC/ClpP-independent manner, and are therefore in need of further optimization. Together, our findings provide a foundation for the development of new tools for protease deregulation as potential antibiotic strategy.

## Results

### High-throughput screen identifies small-molecule activators of *S. aureus* ClpC

To identify compounds that activate the *S. aureus* ClpC/ClpP protease, we conducted a high-throughput screen using a chemically diverse library of approximately 110,000 small molecules. Compounds were screened at a single concentration (40 µM) for increasing ClpC/ClpP proteolytic activity, which was measured using fluorescently labeled FITC-casein as model substrate. Degradation of FITC-casein causes an increase in fluorescence intensity, which was measured in an endpoint experiment after 60 minutes of incubation in the absence and presence of compounds (Figure 1A). On each assay plate, columns 1 + 2 contained ClpC and ClpP as negative controls, and columns 23 + 24 served as a positive control with ClpC, ClpP, and the adapter protein MecA. This enabled normalization for each plate and monitoring of assay performance with Zʹ-factors. Assay quality, as assessed by the Zʹ-factor, ranged between 0 and 0.6 across screening plates, indicating moderate robustness but sufficient separation between signal and background to support hit identification (13 out of 337 assay plates were not taken into account as the Zʹ-values were below zero). The primary analysis, in which we applied a threshold of at least 85% activation relative to the positive control, yielded approximately 900 preliminary hits, corresponding to a relatively high hit rate. To prioritize promising scaffolds and reduce redundancy, hit compounds were analyzed using chemical structure clustering. Only compound families with multiple active analogs were selected for follow-up testing, while singletons (more than 670 compounds) were deprioritized. Representative compounds from prioritized clusters were subsequently selected. Taking into account availabilities, 57 compounds were bought from the vendors from which the compounds were originally acquired for further biochemical and mechanistic characterization.

**Figure 1.**
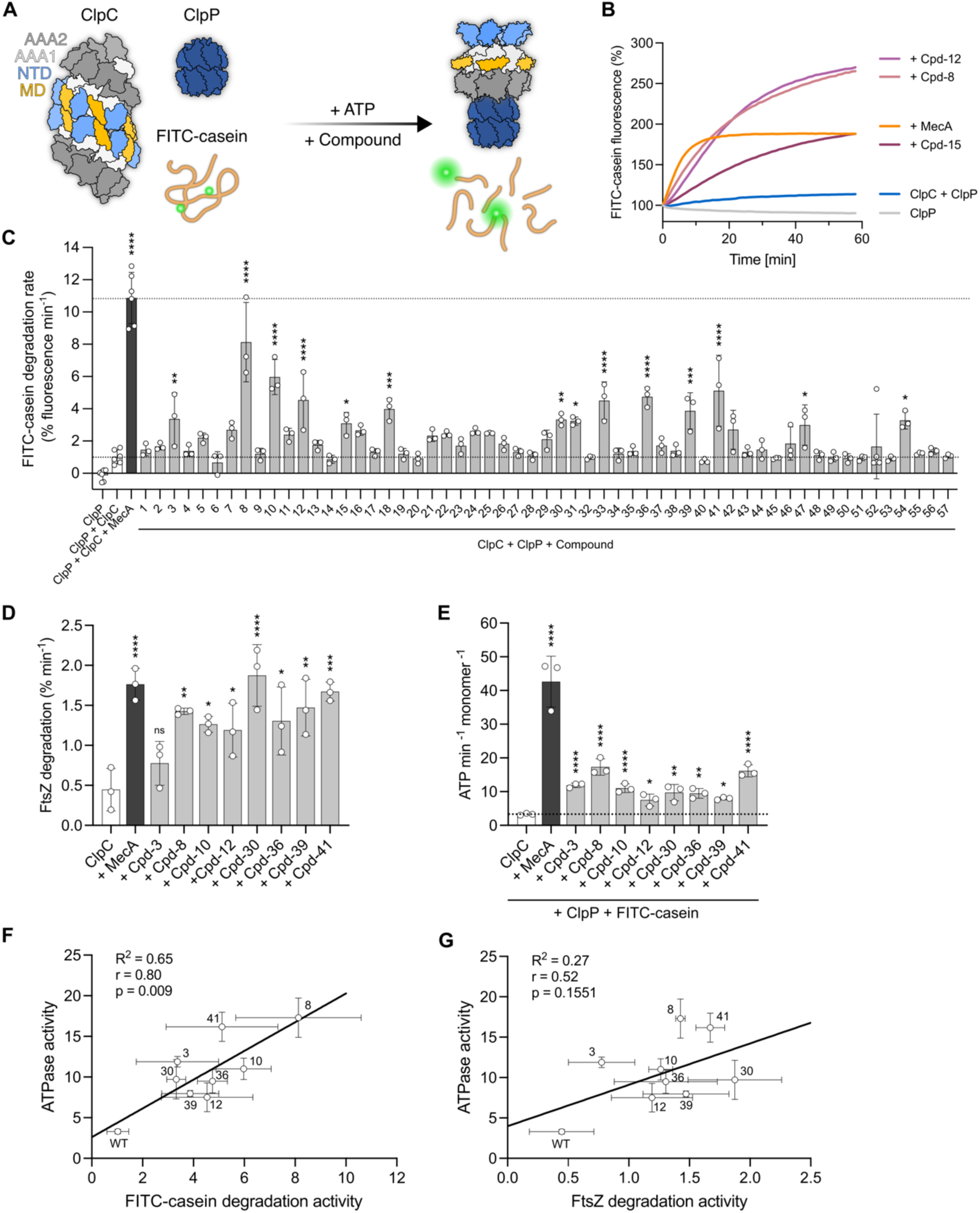
Small molecule activators identified in a high-throughput screen activate *S. aureus* ClpC *in vitro*. (A) Model illustrating ClpC activity control and the functional principle of the FITC-casein degradation assay used for high-throughput screening. In its resting state, ClpC is inactive and unable to engage ClpP. Compound-induced activation promotes hexamer assembly, allowing formation of an active ClpC/ClpP protease that degrades FITC-casein, monitored as fluorescence dequenching. (B) Representative curves of proteolytic activities of ClpP and ClpC + ClpP ± MecA or selected compounds towards FITC-casein. Initial FITC-casein fluorescence was set to 100%. (C) FITC-casein degradation activities were determined in the presence of ClpP and ClpC + ClpP ± MecA or respective compounds (50 µM). Data are represented as mean ±SD, n ≥ 3. (D) FtsZ degradation activities were determined in the presence of ClpP ± MecA or respective compounds (50 µM). Data are represented as mean ±SD, n = 3. (E) ATPase activities of ClpC were determined in the presence of ClpP and FITC-casein ± MecA or respective compounds (50 µM). Data are represented as mean ±SD, n = 3. (F-G) Correlation of ATPase and FITC-casein (F) or FtsZ degradation activities (G) for the respective compounds was calculated using two-tailed Pearson correlation tests. ns = not significant, \**p* < 0.05, \*\**p* < 0.01, \*\*\**p* < 0.001, \*\*\*\**p* < 0.0001 by one-way ANOVA with Dunnett’s post-hoc test (C, D, and E). Positive controls were assessed separately.

### Biochemical validation and characterization of primary hits

Next, the 57 primary hits (Figure S1) were tested in a time-resolved FITC-casein degradation assay for better quantification of their stimulatory impact but also to exclude false-positive candidates (Figures 1B, 1C, and S2A). Approximately 50% of the compounds failed to reproducibly stimulate ClpC/ClpP activity and were classified as false positives. The remaining compounds induced robust FITC-casein degradation, as shown by representative kinetic traces in Figure 1B and calculated degradation rates (% fluorescence increase min^-1^) in Figure 1C. Basal ClpC/ClpP activity was low (1.03% fluorescence increase min^-1^) and increased strongly upon addition of the adaptor protein MecA, which served as a positive control and exhibited the highest initial degradation rate (10.87% fluorescence increase min^-1^). Degradation kinetics of the positive control reached an early plateau as MecA targets itself for degradation by ClpC/ClpP (Figure 1B). In contrast, the small-molecule activators supported sustained protease activity and resulted in higher total FITC-casein degradation after 60 minutes, indicating that they were not consumed during the reaction (Figure S2A). The most significant increase was observed with compound 8 (Cpd-8), which enhanced ClpC/ClpP activity by a factor of 7.9 (8.13% fluorescence increase min^-1^), approaching the degradation rates mediated by MecA (10.6-fold increase). Based on our results, we selected 12 compounds and measured their efficiencies using dose-response curves. The compounds’ potencies were in the low- to moderate micromolar range, with compounds 8, 10, and 41 showing the lowest EC50 values (Cpd-8: 12 µM; Cpd-10: 4.4 µM; Cpd-41: 8 µM) (Figures S2B and S2C). Building on these findings, we excluded Cpd-15, Cpd-18, and Cpd-54 from further analyses and focused on the remaining eight compounds (Cpd-3, Cpd-8, Cpd-10, Cpd-12, Cpd-30, Cpd-36, Cpd-39, and Cpd-41). Lastly, we conducted a ClpP counter-assay to rule out ClpC-independent FITC-casein degradation by ClpP only in the presence of compounds. ClpP alone did not degrade FITC-casein, yet binding of ADEP1, which served as a positive control, caused substrate proteolysis (2.0% fluorescence increase min^-1^) (Figure S2D). None of the tested compounds enabled ClpP to degrade FITC-casein, confirming that they target ClpC.

Next, we used the bacterial cell division protein FtsZ as an alternative, native substrate of ClpC/ClpP (Feng et al. 2013) in an SDS-PAGE-based degradation assay. All eight compounds increased FtsZ degradation rates by 1.7-4.2-fold, with Cpd-30 showing the greatest effect and resulting in total FtsZ degradation up to 100% after 120 min (Figures 1D, S2E and S2F). These results demonstrate that the compound efficacies are largely independent of the substrate. To directly link enhanced proteolytic activity to ClpC activation, we measured the ATPase activity of ClpC in the presence of compounds with or without the addition of either ClpP, FITC-casein, or both (Figures 1E, S2G and S2H). All compounds significantly stimulated ClpC ATPase activity (2.2-4.9-fold), but to a lesser extent than MecA-mediated activation (12.9-fold) (Figure 1E). The most pronounced effects were typically observed in the presence of both ClpP and FITC-casein (Figure S2G), with Cpd-8 and Cpd-41 inducing the highest ATPase activity (5.2-and 4.9-fold, respectively); however, the impact of ClpP and FITC-casein varied across compounds (Figures S2G and S2H). Importantly, we observed a strong correlation between ClpC ATPase activation and ClpC/ClpP proteolytic activities, indicating that compounds act by overriding ClpC activity control (Figures 1F and 1G). Surprisingly, Cpd-10 inhibited ClpC ATPase activity in the absence of ClpP and FITC-casein. This inhibition was countered by adding ClpP, whereas an activating effect was achieved by adding FITC-casein (Figures S2G and S2H). Collectively, these data demonstrate that the identified compounds activate ClpC independently of substrate identity or assay format. Together, these results identify a set of small-molecule activators that deregulate *S. aureus* ClpC, leading to increased ATP turnover and persistent ClpC/ClpP proteolytic activity.

### Compounds trigger ClpC conformational rearrangements that promote ClpP interaction

The increased activity state of ClpC indicates that the compounds influence its assembly, causing hexamer formation and subsequent association with ClpP. Thus, we used analytical size-exclusion chromatography (SEC) to observe ClpC association with ClpP. Here, we used ATPase-deficient ClpC-E280A/E618A (ClpC-DWB) to stabilize potential complexes with ClpP. ClpC-DWB eluted prior to a 670 kDa standard and was unable to form complexes with ClpP, indicating resting state formation (Figure 2A). The addition of MecA induced hexamer formation, enabling efficient protease complex formation as shown by the co-elution of ClpC-DWB, MecA, and ClpP prior to the ClpC resting state. Presence of Cpd-8, Cpd-10, or Cpd-41 also induced the formation of ClpC/ClpP assemblies, consistent with ClpC/ClpP activation. Complex formation was, however, less efficient, which could be explained by dissociation of the compounds during the SEC runs. Furthermore, compound-triggered ClpC/ClpP complexes were larger than the canonical ClpC/ClpP/MecA complex, eluting close to the void volume of the SEC column. To understand the structural organization of these compound-induced ClpC/ClpP assemblies, we analyzed the early-eluting SEC fractions by negative-stain electron microscopy (EM) (Figure 2B). ClpC/ClpP complexes appeared very large and highly heterogeneous, yet consisted of interacting ClpC hexamers interconnected by ClpP (Figure 2B, Figure S3). Exemplarily, we obtained 2D class averages of some complexes in the presence of Cpd-8, which we suggest represent building blocks of larger assemblies (Figures 2B and S3A). The simplest element consisted of two ClpC hexamers interacting head-to-head, presumably through their coiled-coil middle domains (MDs), while being associated with ClpP. Such ring-dimer formation has recently been shown for the AAA+ disaggregase ClpL (Bohl et al. 2024; Kim et al. 2020) and for ClpP-bound *S. aureus* ΔN-ClpC, lacking its N-terminal domain, or partially in the presence of the substrate FITC-casein and phosphoarginine (pArg) acting as allosteric activator (Jenne et al. 2025). Other building blocks consisted of four ClpC hexamers forming a tetrahedral assembly that associates with multiple ClpPs (Figure 2B). ClpC tetrahedrons have been shown to form in the presence of the toxic cyclic peptide Cyclomarin A (Maurer et al. 2019; Taylor et al. 2022), pArg-substrates (Jenne et al. 2025; Morreale et al. 2022) or derepressed ClpC mutants in the presence of substrate FITC-casein (Jenne et al. 2025), respectively. In such a tetrahedral assembly, ClpC hexamers interact again via head-to-head interacting MDs. We also identified complexes comprising at least five ClpC hexamers, an assembly that has not been reported yet. Similar organizations of large ClpC/ClpP complexes were observed in the presence of Cpd-10 and Cpd-41 (Figures S3B and S3C). Taken together, the tested compounds induce ClpC hexamer formation and the subsequent development of large ClpC assemblies with remarkably high structural variability. These assemblies likely form via MD-MD interactions and have ClpC AAA2 rings accessible for ClpP docking. This, combined with ClpP’s ability to bind two ClpC hexamers, allows for the formation of extensive networks of ClpC assemblies interconnected by ClpP.

**Figure 2.**
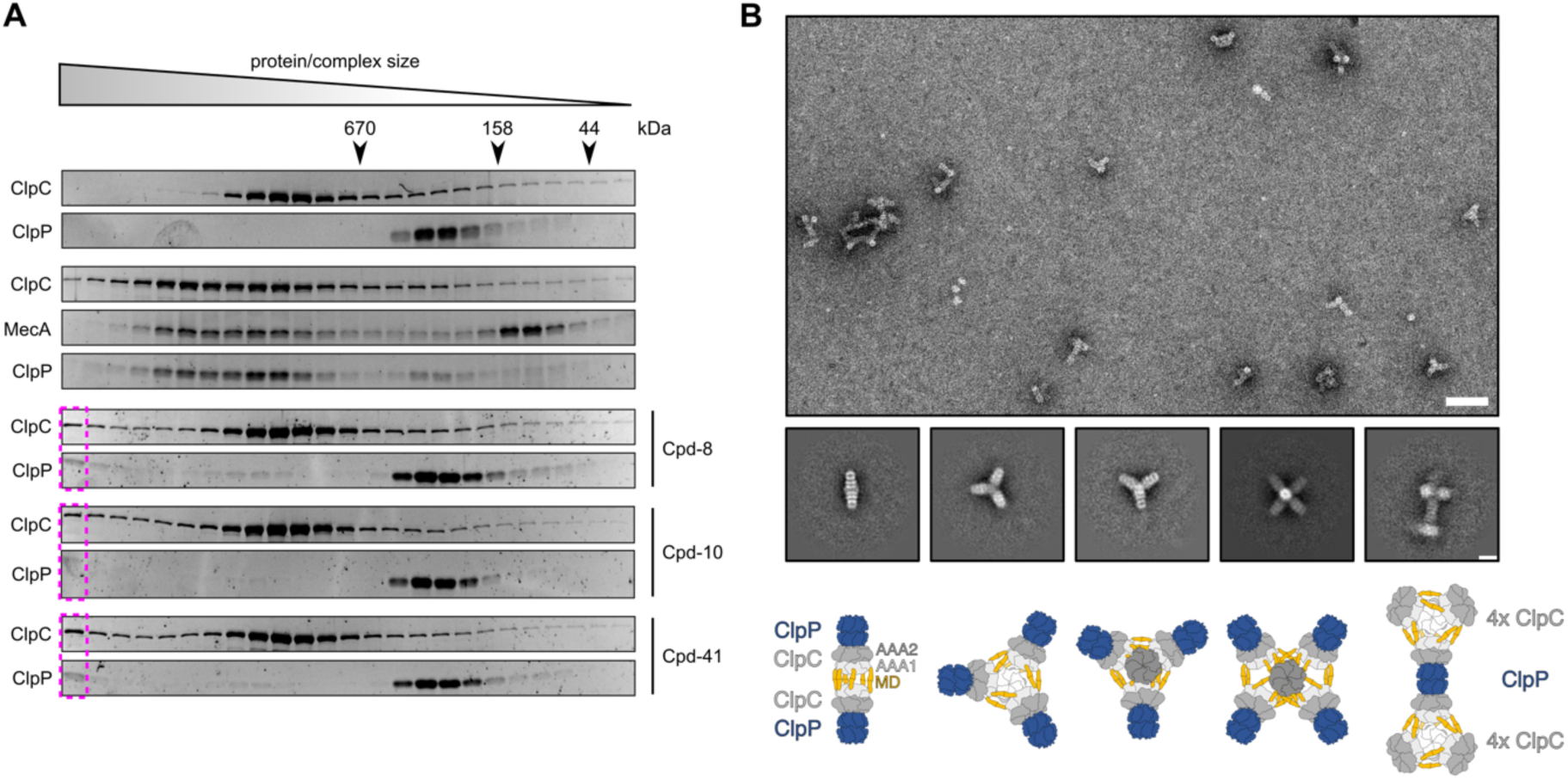
Compounds trigger conformational rearrangements that activate ClpC and promote ClpP interaction. (A) Complex formation of ATPase-deficient ClpC-E280A/E618A (DWB) with ClpP and FITC-casein ± MecA or respective compounds (50 µM) was monitored by Superose 6 size-exclusion chromatography. Elution fractions were analyzed by SDS-PAGE and SyproRuby staining. (B) Negative stain EM of ClpC-DWB/ClpP complexes formed in the presence of FITC-casein and Cpd-8 as isolated from SEC runs (see magenta boxes in (A)). Representative micrograph (scale bar = 50 nm) and 2D class averages (scale bar = 20 nm) reveal the formation of variable ClpC-DWB/ClpP complexes including rods, tetrahedrons, and other higher-order complexes composed of ClpC-DWB hexameric rings that interact head-to-head through coiled-coil M-domains. These complexes are connected through tetradecameric ClpP, which offers two docking sites for ClpC hexamers. The structural organizations of representative 2D classes are depicted as cartoons highlighting ClpP (dark blue) and ClpC domains (MD = orange, AAA1 = white/light grey, AAA2 = grey).

### The N-terminal domain (NTD) serves as compound binding site

Next, we aimed to identify the compound binding sites to better understand their mechanisms of action (MoA). The N-terminal domain of Mtb ClpC1 is the target of all currently described toxic cyclic peptides, which deregulate ClpC1 and cause severe toxicity (Vasudevan et al. 2013; Wolf et al. 2019; Jagdev et al. 2023; Wolf et al. 2020). Furthermore, the NTD has recently been identified as a central regulatory allosteric hub in ClpC activity control, stabilizing the resting state and integrating various input signals like adaptor proteins (e.g., MecA) or the pArg-degron tag for ClpC activation (Jenne et al. 2025). Therefore, we first used a ClpC variant lacking its NTD (ΔN-ClpC) and measured its ATPase activity in the absence and presence of compounds (Figure 3A). None of the compounds stimulated ΔN-ClpC ATPase activity, suggesting that the NTD is the compound-binding site. Cpd-10 even inhibited ΔN-ClpC ATPase activity by a factor of 3.3, while the addition of FITC-casein and ClpP lessened this inhibitory effect. This inhibition, which we also observed in part for ClpC-WT (Figures S2G and S2H), suggests the existence of a secondary Cpd-10-binding site with lower affinity outside the NTD. Next, we evaluated whether compounds could enhance the proteolytic activity of ΔN-ClpC/ClpP. None of the compounds was able to increase proteolytic activity, supporting our assumption that all compounds bind to the NTD (Figure 3B). ΔN-ClpC-F436A, a constitutively activated ClpC variant, served as positive control documenting that FITC-casein degradation is possible without the NTD, which harbors a hydrophobic groove that binds unfolded proteins like casein (Jenne et al. 2025; Rosenzweig et al. 2015). To further document compound binding to the NTD, we used a reciprocal approach and added an excess of isolated NTD in both FITC-casein degradation and ATPase assays to determine whether compounds are sequestered by free NTDs, thereby preventing compound-mediated ClpC-WT activation (Figures 3C and 3D). NTD excess reduced the stimulatory effects of all compounds on FITC-casein degradation activities of ClpC-WT/ClpP to varying degrees (Figure 3C). Furthermore, the increase of ClpC ATPase activities by compounds was completely abolished (Figure 3D). We infer that all eight tested compounds deregulate ClpC by binding to its NTD.

**Figure 3.**
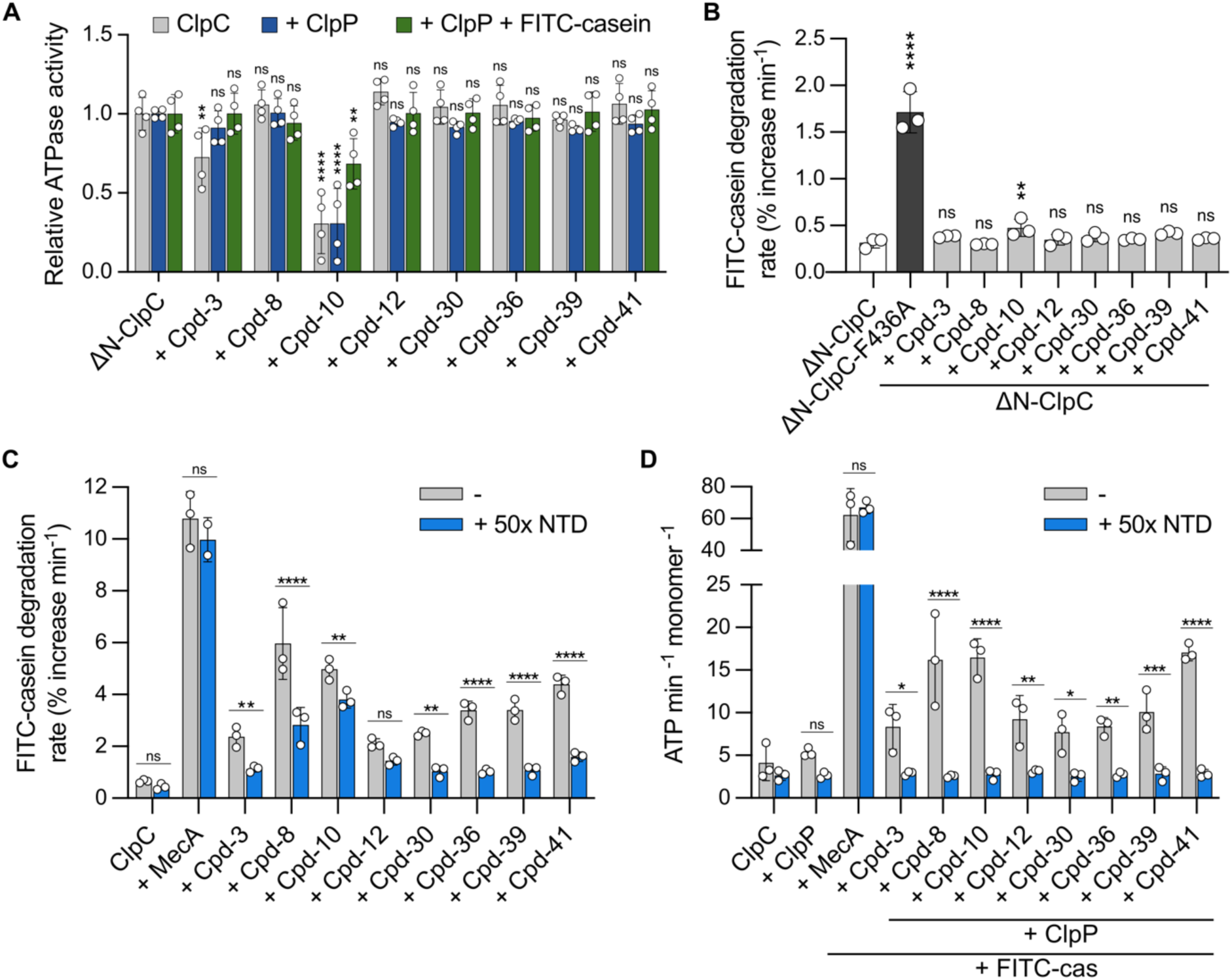
The N-terminal domain (NTD) serves as compound binding site. (A) ATPase activities of ΔN-ClpC alone or in the presence of ClpP or ClpP + FITC-casein were determined ± respective compounds (50 µM). Activities were normalized to ΔN-ClpC alone (set to 1) for each condition. Data are represented as mean ±SD, n = 4. (B) FITC-casein degradation activities of ΔN-ClpC were determined in the presence of ClpP ± respective compounds (50 µM). ΔN-ClpC-F436A was used as a positive control because ΔN-ClpC cannot bind MecA. Data are represented as mean ±SD, n = 3. (C-D) FITC-casein degradation (C) or ATPase activities (D) of ClpC-WT + ClpP + FITC-casein were determined in absence or presence of a 50-fold excess of isolated NTD at the indicated conditions. Compound concentrations = 25 µM. Data are represented as mean ±SD, n = 3. ns = not significant, \**p* < 0.05, \*\**p* < 0.01, \*\*\**p* < 0.001, \*\*\*\**p* < 0.0001 by two-way ANOVA with Dunnett’s post-hoc test (A), one-way ANOVA with Dunnett’s post-hoc test (B), or two-way ANOVA with Holm-Šidák’s post-hoc test (C and D). Positive controls were assessed separately.

### Compounds putatively bind to the hydrophobic groove and the pArg1-site of the NTD

In order to map the compound binding sites on the NTD, we turned to *in silico* docking simulations. First, we used DiffDock-L (Corso et al. 2022, 2024), a generative model for ligand docking, to screen for putative ligand binding sites (Figures 4A, 4B and S4). Strikingly, all compounds were predicted to bind to one of the three known ligand and substrate binding sites of the NTD, namely the hydrophobic groove and both pArg1/pArg2 binding sites, which independently bind phosphorylated arginines (pArg) as a degron (Trentini et al. 2016). Most compounds primarily docked to the pArg1-site, while the hydrophobic groove of the NTD, which is the binding site for all ClpC1-targeting cyclic peptides (Vasudevan et al. 2013; Wolf et al. 2019; Jagdev et al. 2023; Wolf et al. 2020), was typically predicted with lower frequencies (Figure 4B). Building on these findings, we performed HADDOCK simulations (Honorato et al. 2024, 2021) biased toward the hydrophobic groove and the pArg1- and pArg2-sites (Figures 4C and S5). In contrast to DiffDock, HADDOCK relies on a well-established, physics-based energy function and a staged workflow (rigid-body docking, semi-flexible refinement, and final refinement in explicit water), which allows local side-chain/backbone adjustments to produce a more complementary ligand-bound conformation. The resulting HADDOCK score is computed as a weighted linear combination of intermolecular energies (e.g., van der Waals, electrostatics, and desolvation), and is commonly used as a physically interpretable proxy for the relative plausibility of a given pose. Moreover, it was designed to be compatible with ambiguous interaction restraints. These *in silico* predictions generally support binding of most compounds to the hydrophobic groove (Cpd-3, Cpd-8, Cpd-12, Cpd-30, Cpd-39) and strongly oppose binding to the pArg2-site for all compounds based on the total score of the binding poses. However, some predictions showed similar (Cpd-10, Cpd-36) or even better (Cpd-41) average scores for the pArg1-site, qualifying it as an alternative compound binding site. On this basis, we assessed whether the compounds could outcompete the adaptor variant MecA-mEOS for ClpC interaction. MecA-mEOS is composed of the C-terminal domain of MecA, which binds to both ClpC pArg1- and pArg2-sites, and the mEOS fluorophore. MecA-mEOS activates ClpC and is subjected to ClpC/ClpP-mediated degradation (Azinas et al. 2025). We reasoned that compound binding to either pArg1 or pArg-2 site should block MecA-mEOS binding and prevent its degradation. Indeed, Cpd-10 and Cpd-41 effectively abolished MecA-mEOS degradation, indicating they bind to at least one of the pArg-binding sites (Figure 4D). Cpd-8 slightly reduced MecA-mEOS degradation, whereas all other compounds showed no significant inhibition but rather a slight stimulatory effect. To probe for compound binding to the pArg1-site, we used the ClpC mutant T7D, which is located at the pArg1 center (Figure 4E). ClpC-T7D is deregulated and exhibits high autonomous proteolytic activity (Figure S6A) (Jenne et al. 2025). Intriguingly, most compounds still enhanced ClpC-T7D activity, whereas Cpd-10 and Cpd-41 no longer did so (Figure 4F). These experimental findings support HADDOCK simulations of Cpd-10 and Cpd-41, indicating that these compounds activate ClpC by targeting the pArg1-site.

**Figure 4.**
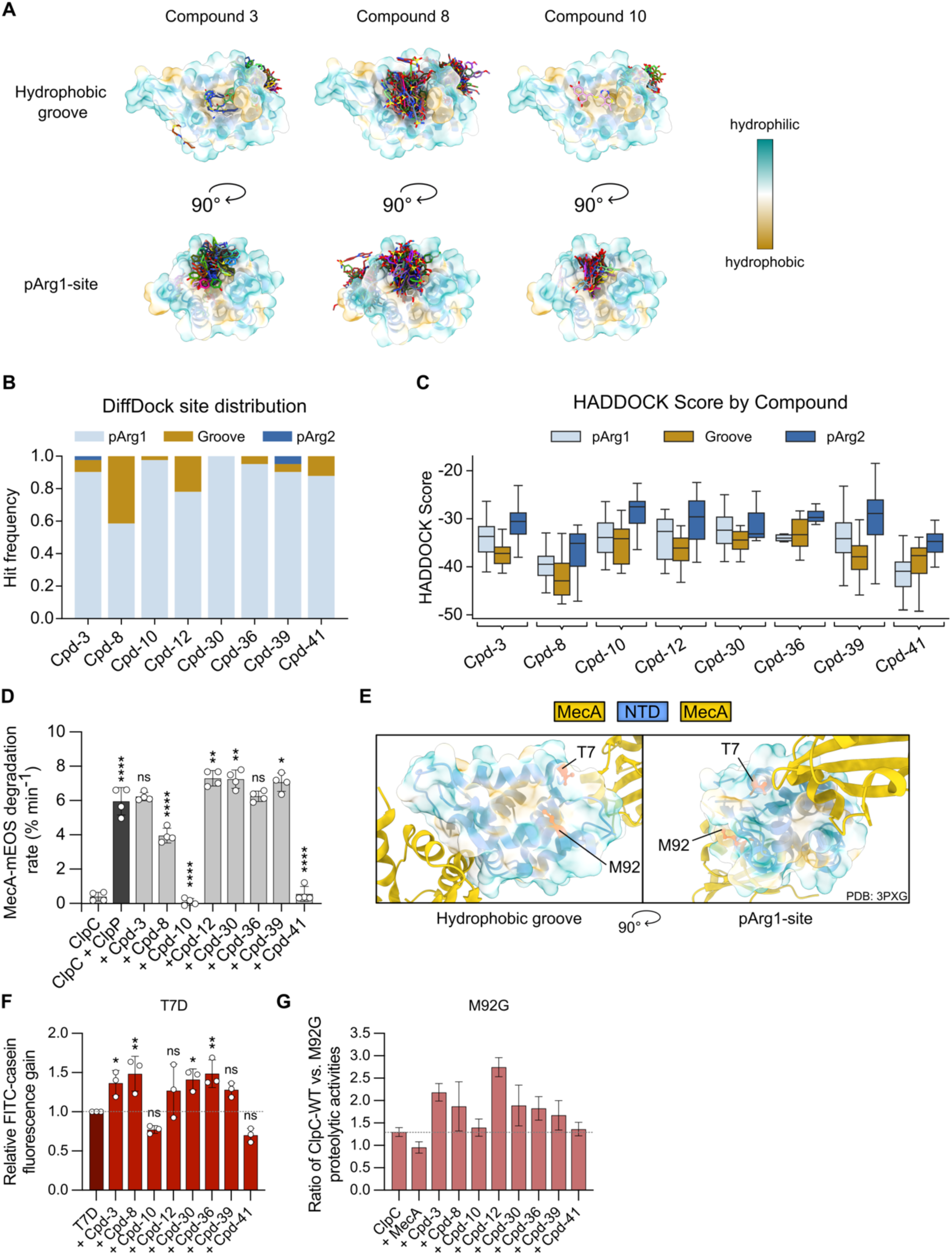
Compounds putatively bind to the hydrophobic groove and pArg1-site of the NTD. (A-B) DiffDock docking simulations (n = 40) of selected compounds to the ClpC N-terminal domain. The NTD is shown as a surface colored by hydrophobicity, two orthogonal views highlighting the hydrophobic groove and the pArg1 site, with all predicted poses displayed for selected compounds (A). Predicted binding poses were clustered by binding site and quantified as the fraction of solutions occupying each site, displayed as stacked bar diagram. (C) Targeted HADDOCK simulations to the hydrophobic groove and both pArg1-/pArg2-sites were performed. Total HADDOCK scores for the corresponding compound docked to each indicated binding site are shown. The solid line in the boxes indicates the median, and the box outlines the 25th and 75th percentiles, respectively. (D) MecA-mEOS degradation by ClpC/ClpP was determined ± respective compounds (50 µM). Data are represented as mean ±SD, n = 4. (E) Structural representation of ClpC in complex with MecA (Wang et al. 2011), highlighting hydrophobicity of the ClpC-NTD and residues subjected to mutagenesis located at the hydrophobic groove (M92) or the pArg1-site (T7) in red. (F) FITC-casein degradation activities of ClpC-T7D were determined ± respective compounds (50 µM) in the presence of ClpP. Data are represented as mean ±SD, n = 3. (G) FITC-casein degradation activities of ClpC-WT and ClpC-M92G were determined ± respective compounds (50 µM) in presence of ClpP. Data represent the factor of the reduced activity upon M92G mutation for each condition (Ratio WT/M92G). Data are represented as mean ±SD, n ≥ 3. SDs have been propagated from the SDs of degradation activities of WT and M92G under the displayed conditions, respectively. ns = not significant, \**p* < 0.05, \*\**p* < 0.01, \*\*\**p* < 0.001, \*\*\*\**p* < 0.0001 by one-way ANOVA with Dunnett’s post-hoc test (C), with Fisher’s LSD (E), ns = not significant, *p* = 0.1662 for Cpd-10, *p* = 0.1494 for Cpd-12, *p* = 0.0955 for Cpd-39, *p* = 0.0803 for Cpd-41; \**p* = 0.0394 for Cpd-3, * *p* = 0.0173 for Cpd-30; \*\**p* = 0.0061 for Cpd-8, ** *p* = 0.0079 for Cpd-36. Positive controls were assessed separately.

To clarify the binding sites of the other six compounds, we aimed to mutate the NTD’s hydrophobic groove. Several studies have reported that mutations in the hydrophobic groove of the homologous NTD of Hsp104 strongly decrease domain stability and are thus unsuitable for analysis (Sweeny et al. 2020; J. Lee et al. 2017; Rosenzweig et al. 2015). Therefore, we followed a successful approach previously reported for the NTD of Hsp104 and designed the double-mutant ClpC-L14D-I88R to create a salt bridge, which blocks access to the groove (Harari et al. 2022) (Figure S6B). However, this mutant could no longer interact with MecA, as evidenced by ATPase and MecA-mEOS degradation assays (Figures S6C and S6D). The mutated sites are not involved in MecA binding, pointing to structural defects. We additionally introduced these mutations into the isolated NTD and determined its melting temperature using nano differential scanning fluorimetry (nanoDSF). The melting temperature massively dropped from 63°C (WT) to 38°C (L14D-I88R) upon NTD mutation (Figure S6E), confirming impaired structural integrity. Hence, we chose Met92 for mutagenesis, which is located at the edge of the hydrophobic groove and has its side chain pointing toward the groove’s center (Figure 4E). FITC-casein degradation activities of ClpC-WT and ClpC-M92G in the presence of MecA were similar, indicating that the NTD remains structurally intact upon mutation (Figure S6F). Stimulation of ClpC-M92G proteolytic activity by Cpd-10 and Cpd-41 was comparable to ClpC-WT (Figure 4G), in accordance with previous results suggesting that they bind to the pArg1-site. Enhancement of ClpC-M92G activity by all other compounds (Cpd-3, Cpd-8, Cpd-12, Cpd-30, Cpd-36, Cpd-39) was reduced as compared to ClpC-WT (1.67-2.75-fold), supporting binding of these six compounds to or near the hydrophobic groove in agreement with HADDOCK simulations (Figure 4G). We conclude that compounds target two distinct sites in the ClpC NTD: the hydrophobic groove and the pArg1 site.

### Dissection of Cpd-8 binding mode and its potential mechanism of action

To directly determine how the most potent compounds Cpd-8, Cpd-10, and Cpd-41 bind to the ClpC-NTD, we conducted co-crystallization experiments. We could obtain crystal structures of ClpC-NTD with bound Cpd-8 at 1.90 Å and in the apo state at 1.78 Å resolution (Table S1). Interestingly, both data sets have the same space group P4_3_2_1_2 with very similar cell dimensions and one protein molecule per asymmetric unit. Cpd-8 binds to a part of the hydrophobic groove, and the entire ligand shows clear electron density (Figures 5A, 5B and S7A). A closer look at the binding site reveals that only a part of Cpd-8, dubbed as Cpd-8a (1-(3,5-bis(4-methoxyphenyl)-4,5-dihydro-1H-pyrazol-1-yl)ethan-1-one), forms protein-ligand contacts (Figures S7B-D). The other part of the ligand, referred to as Cpd-8b (4-(4-oxo-4,5-dihydro-1H-pyrazolo[3,4-d]pyrimidin-1-yl)benzenesulfonamide), is involved in a crystal contact with a symmetry-equivalent ligand molecule. Notably, protein and ligand form only one hydrogen bond, namely between Gly4 and the carbonyl group of Cpd-8a, whereas all other protein-ligand interactions involve hydrophobic contacts (Figures S7B and S7C). The superposition of the apo and complex structure (Figure S7E) shows a rearrangement of the first three N-terminal residues upon ligand binding. These three residues cover one of the two Cpd-8a methoxyphenyl moieties. Significantly smaller movements can be observed for the Met92 Cε and Tyr80 sidechain. Ligand binding does not induce any other large movements in ClpC-NTD, which is indicated by a low rmsd(C_α_) of 0.14 Å for the superposition of residues 4 to 144 of apo and complex structure.

**Figure 5.**
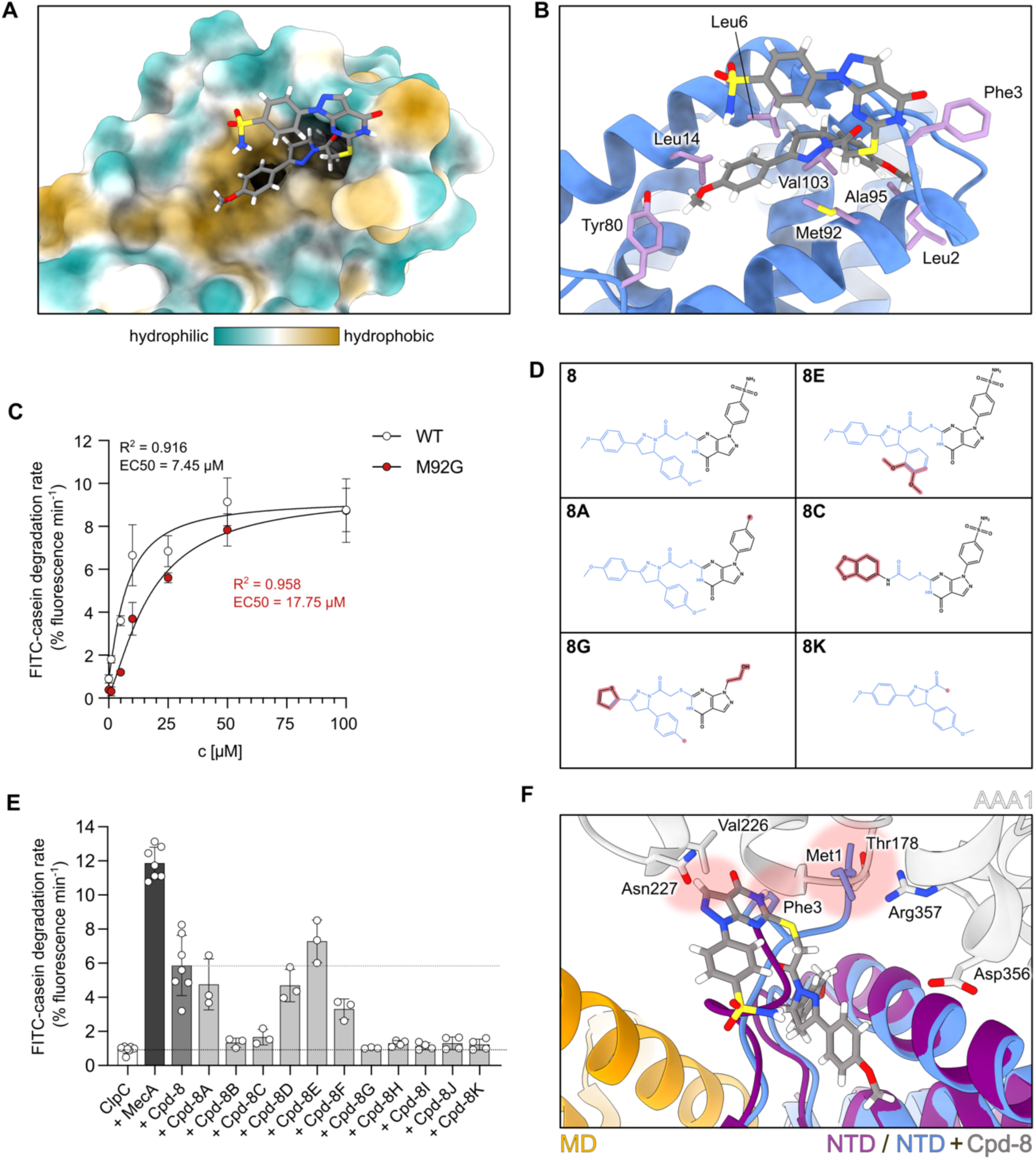
Dissection of Cpd-8 binding mode and its potential mechanism of action. (A-B) Crystal structure of ClpC-NTD with bound Cpd-8 (carbon atoms in grey) in surface representation colored by hydrophobicity (A) and as cartoon representation (blue) showing selected residues (plum) involved in protein-ligand contacts (B). (C) Dose-response curves of ClpC-WT and ClpC-M92G were determined by monitoring FITC-casein degradation activities at increasing Cpd-8 concentrations in presence of ClpP. EC50 values were obtained by fitting agonist–response data with a four-parameter variable-slope nonlinear regression model. Data are represented as mean ±SD, n = 4. (D) Chemical structures of Cpd-8 and selected derivatives. Structural elements of Cpd-8 involved in protein–ligand contacts are displayed in blue. Modifications of respective derivatives are highlighted in red. (E) FITC-casein degradation activities of ClpC ± Cpd-8 and respective derivatives (50 µM) were determined in presence of ClpP. Data are represented as mean ±SD, n ≥ 3. (F) The crystal structure of the ClpC N-terminal domain (blue) with bound Cpd-8 (carbon atoms in grey) was superimposed on *S. aureus* ClpC in resting state (PDB: 9QCL), AAA1: grey, MD: orange, NTD: purple. Areas with potential steric clashes (VDW overlaps ≥ 0.6 Å) between the resting state and the N-terminus of ClpC-NTD and bound Cpd-8 are marked in light red.

To biochemically validate the binding position of Cpd-8 observed in the crystal structure, we used the previously described ClpC-M92G mutant, as this residue makes multiple contacts with Cpd-8 (Figures 5B, S7B and S7C), and determined dose-response curves. The EC50 increased by a factor of 2.4 with the M92G mutation (Figure 5C), indicating decreased affinity in agreement with the determined binding mode.

To assess the functional relevance and evaluate the structure-activity relationship (SAR) of Cpd-8, we tested the efficiencies of commercially available derivatives of Cpd-8 (Figures 5D and S8A) in FITC-casein degradation assays. Modifying one 3-methoxybenzene moiety of Cpd-8a to 1,2-dimethoxybenzene (Cpd-8E) resulted in a slight increase in FITC-casein degradation activity compared to Cpd-8 (Figure 5E). However, more extensive modifications of Cpd-8a generally caused a complete loss of the ability to enhance FITC-casein degradation activities (Cpd-8B, Cpd-8C, Cpd-8G, Cpd-8I). This underlines the central role of Cpd-8a for NTD binding. However, the fragment Cpd-8a alone (Cpd-8K) is insufficient for ClpC activation, indicating that parts of the second moiety, Cpd-8b, also play a crucial role. Substitution of the sulfonamide group with fluoride (Cpd-8A) in the Cpd-8b fragment caused only a slight decrease in proteolytic activity, implying negligible functionality. Replacing Cpd-8b with another large moiety (Cpd-8H) no longer allowed for ClpC activation, indicating that the Cpd-8b fragment cannot simply be swapped out for another bulky group. Thus, we infer that Cpd-8b also contributes to ClpC activation, yet which part of it remains unresolved.

The toxic cyclic peptides cyclomarin A, lassomycin, rufomycin, and ecumicin, which deregulate Mtb ClpC1, bind directly to the center of the hydrophobic groove, making contact with the highly conserved residue Lys85 (Vasudevan et al. 2013; Wolf et al. 2019; Jagdev et al. 2023; Wolf et al. 2020). This residue forms a salt bridge with Asp356 in the AAA1 domain, which is crucial for resting state formation and activity control, and has been proposed as the reason for ClpC1 deregulation by cyclic peptides (Jenne et al. 2025). The binding site of Cpd-8 is slightly different from the previously described cyclic peptides and does not make contact with Lys85 (Figure S8B). We speculate that re-orientation of the ClpC N-terminus could clash with the resting state and destabilize it, leading to ClpC activation. Indeed, superimposing the co-crystal structure with the cryo-EM structure of the ClpC resting state (PDB: 9QCL) revealed serious steric clashes of the N-terminus and the pyrazolo-pyrimidonone moiety of Cpd-8b with the AAA1 domain (Figure 5F), offering a potential mechanistic explanation of how Cpd-8 activates ClpC.

In summary, using co-crystallization, mutagenesis and compound derivatives, we identified and validated the Cpd-8 binding site, established key structure–activity relationships, and propose a possible mechanism of action.

### Dissection of Cpd-10 binding mode and its potential mechanism of action

HADDOCK simulations and experimental evidence support binding of Cpd-10 to the conserved pArg1-site. This regulatory pocket is particularly interesting because its druggability has not been demonstrated for small molecules so far. Our investigation of Cpd-10 concentrated on the predicted binding conformations from the earlier HADDOCK simulations (Figures 4C and S5), specifically examining the binding poses of the top two docking clusters, 1 and 7. The orientation of the aesculetin moiety (hereafter referred to as Cpd-10a) varied to some degree between the two clusters, while the second half of Cpd-10, 5,6-dimethylthieno[2,3-d]pyrimidine (hereafter referred to as Cpd-10b), interacted with the NTD in a similar manner (Figure 6A). In cluster 1, Cpd-10 forms four hydrogen bonds with the NTD, while in cluster 7, only one hydrogen bond is predicted. In contrast to Cpd-8, both parts of Cpd-10 directly interact with the pArg1-site. Superimposition of the cryo-EM structure of the ClpC resting state with Cpd-10-bound NTD revealed significant steric clashes with the tip of the regulatory MDs (Figure 6B). These clashes involve methyl groups of Cpd-10b with Ala429 and Ala430 in both clusters, and Cpd-10a with Glu426, with the clashes being more pronounced in cluster 1. Glu426 has been linked to the regulation of ClpC activity through stabilizing the resting state via NTD interactions. When mutated (ClpC-E426A), this stabilization is lost, leading to destabilization of the resting state and autonomous ClpC/ClpP proteolytic activity (Jenne et al. 2025; Carroni et al. 2017). We therefore propose that Cpd-10, when bound to the NTD, will clash with the MD tip, triggering ClpC activation.

**Figure 6.**
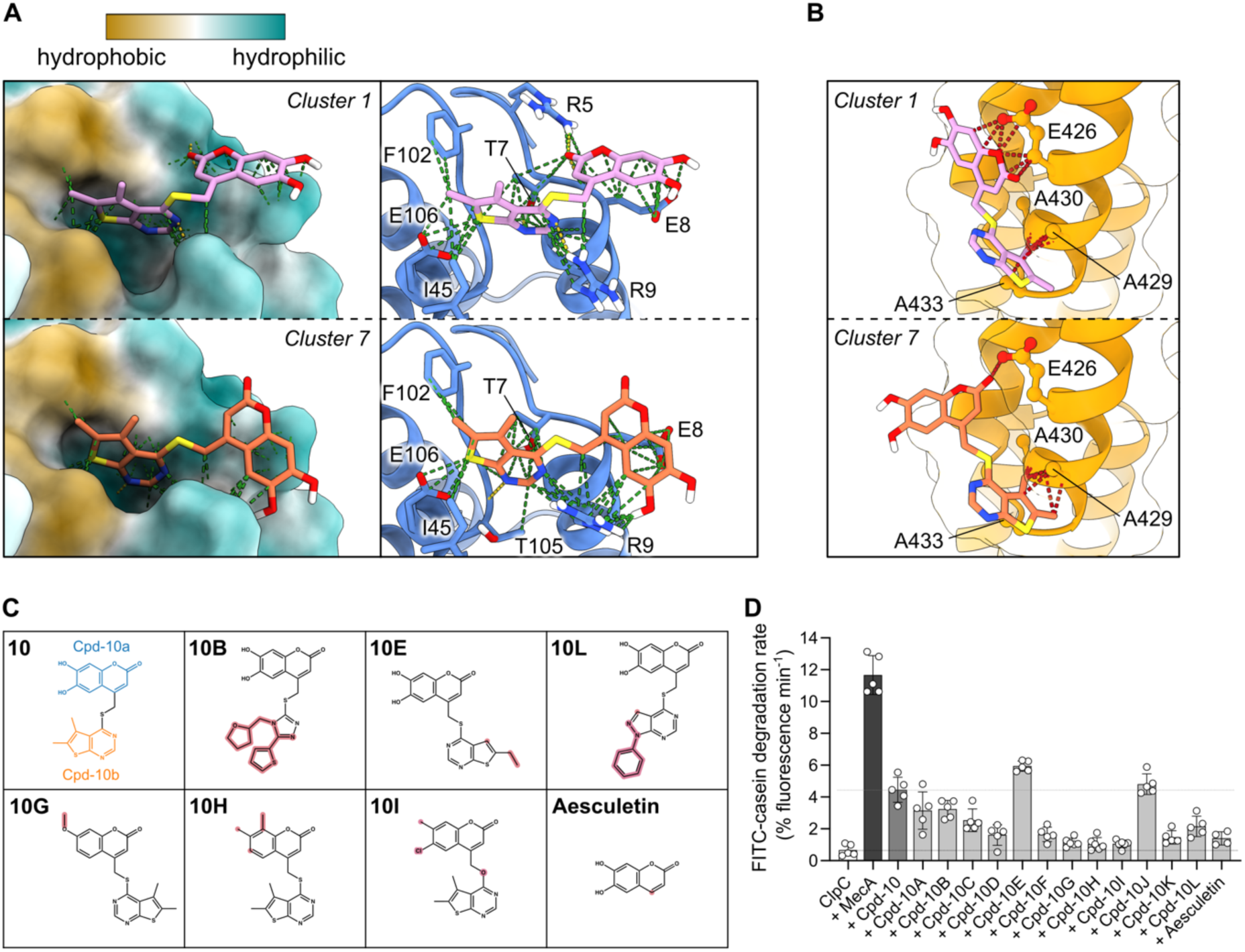
Dissection of Cpd-10 binding mode and its potential mechanism of action (MoA). (A) The top two HADDOCK docking simulations (clusters 1 and 7) of Cpd-10 binding to the NTD are illustrated: a surface representation colored by hydrophobicity (left) and a cartoon view showing hydrogen bonds and contacts (yellow and green dotted lines, respectively, on the right). (B) Superimposition of the cryo-EM structure of the ClpC resting state (Jenne et al. 2025) with the predicted binding poses of Cpd-10 from clusters 1 and 7. Steric clashes between Cpd-10 and the tip of the regulatory middle domains, including involved residues, are indicated. (C) Chemical structures of Cpd-10 and selected derivatives. Structural elements of Cpd-10 are displayed in blue (Cpd-10a) and orange (Cpd-10b), respectively. Modifications of respective derivatives are highlighted in red. (D) FITC-casein degradation activities of ClpC ± Cpd-10 and respective derivatives (50 µM) were determined in the presence of ClpP. Data are represented as mean ±SD, n ≥ 4.

To assess whether Cpd-10’s predicted binding poses align with experimental structure–activity relationships, we examined several commercially available derivatives (Figures 6C and S9) using FITC-casein degradation assays and evaluated their activities in the context of the described top two docking clusters 1 and 7. Even slight modifications of Cpd-10a significantly impair or eliminate the derivatives’ ability to activate ClpC 3.9-9.9-fold, such as Cpd-10G, Cpd-10D, Cpd-10F, Cpd-10H, and Cpd-10I (Figure 6D, Figure S9), highlighting its crucial role in NTD binding. However, Cpd-10a (Aesculetin) alone does not suffice for ClpC activation (Figure 6D), emphasizing the importance of the Cpd-10b moiety for functionality. Although even extensive substitutions of Cpd-10b—such as Cpd-10J, Cpd-10A, Cpd-10C, and Cpd-10B—are generally well tolerated, they cannot be entirely arbitrary. This is evidenced by the decrease in ClpC activity when Cpd-10L and Cpd-10K are present, with reductions of 2.7-fold and 4.1-fold, respectively. On the contrary, removal of the 5-methyl group from the thiophene ring of Cpd-10b and extension of the 6-methyl group to a 6-ethyl group (Cpd-10E) slightly improved compound efficiency (Figure 6D). This shows that Cpd-10b is indeed functionally relevant and needs to satisfy specific criteria.

In light of the SAR results, we assessed the accuracy of HADDOCK clusters 1 and 7, taking into account how respective derivative modifications could affect NTD-binding and affinity, along with potential clashes with the ClpC resting state that could trigger ClpC deregulation. Overall, cluster 1 is strongly favored by the observed effects of Cpd-10 derivatives on FITC-casein degradation activities as the effects of the respective modifications generally correspond well to expected changes in the context of this docking pose. On the contrary, cluster 7 is incompatible with the observed effects for the modifications seen for Cpd-10J, Cpd-10B and Cpd-10L, as the added or modified (hetero-) aromates would clash extremely with the NTD, yet these derivatives still at least partially activate ClpC/ClpP. Collectively, the SAR trends therefore support cluster 1 as the most consistent binding model.

### Compounds induce ClpC/ClpP independent toxicity in *Staphylococcus aureus*

To examine the physiological effects of the compounds as growth inhibitors of S*taphylococcus aureus*, we monitored growth curves at 37°C using a WT strain (ATCC 25923) and subsequently assessed cell viability through spot dilution series (Figures 7A, 7B and S10A). All eight tested compounds reduced cell growth and delayed entry into the logarithmic growth phase at high concentrations (200 µM) (Figure 7A). In particular, Cpd-10 fully prevented growth, similar to the well-known antibiotic vancomycin. Cell viability decreased by 1.8-3.7 times across all compounds, except for Cpd-10, which led to a 90-fold reduction (Figures 7B and S10A). To document that growth impairment is caused by deregulation of ClpC/ClpP, we tested all compounds on two previously characterized knockout strains (NCTC 8325-4): *clpC::erm*, which has both AAA+ domains deleted through a long in-frame deletion (Frees et al. 2004), and *ΔclpP* (Frees et al. 2003). Unfortunately, the growth of *clpC::erm* and *ΔclpP* was still impaired by compounds, and the inhibitory effects were even worse than observed for the WT strain (Figure 7C). This demonstrates that the toxicity observed across all compounds does not depend on ClpC/ClpP, but rather likely results from effects of unknown origin, which become enhanced in the absence of ClpC/ClpP (Figure 7C). Similar results were obtained at low compound concentrations (10/50 µM), and ClpC-independent growth inhibition was observed for all compounds except Cpd-12 (Figure S10B).

**Figure 7.**
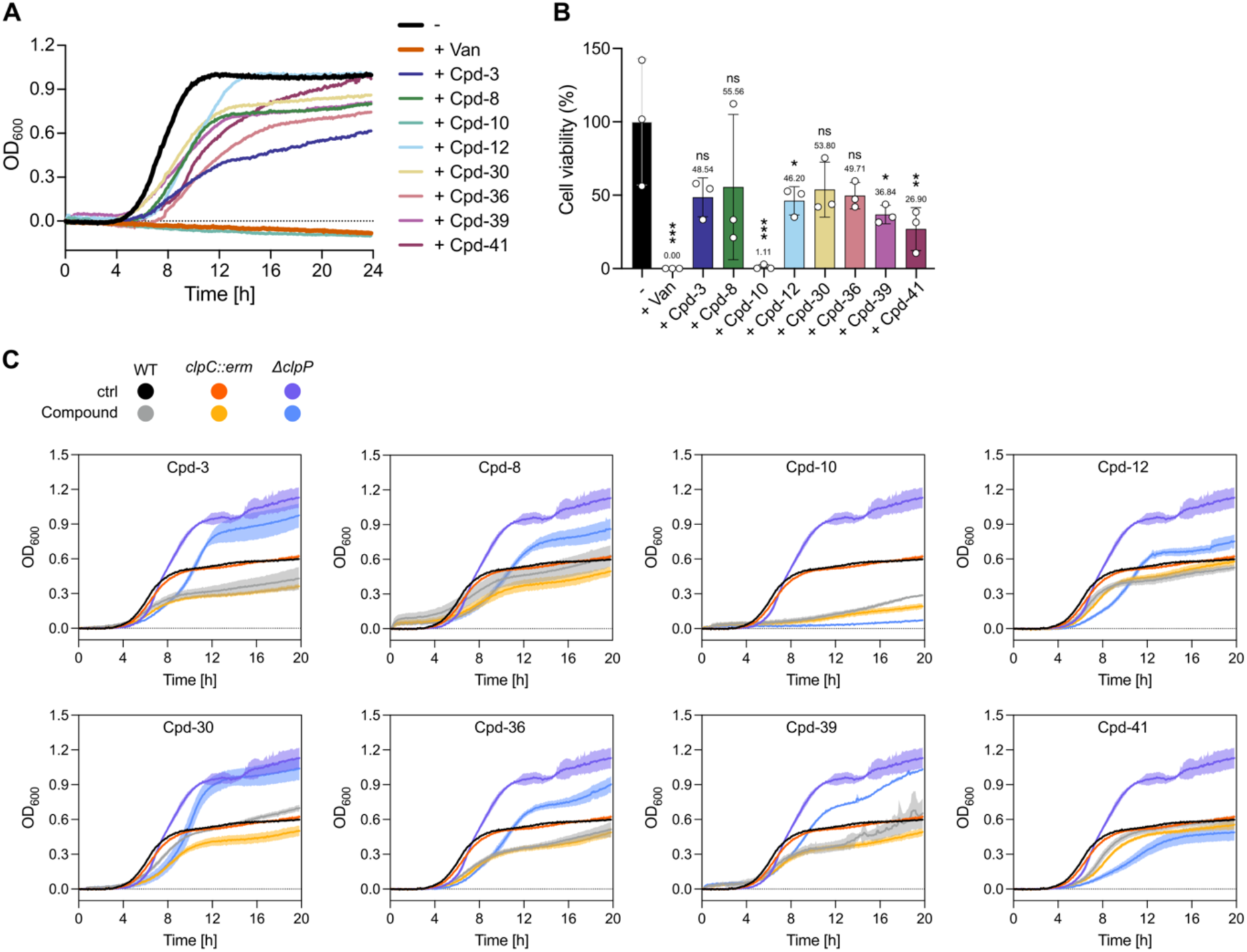
Compounds induce ClpC/ClpP independent toxicity in *Staphylococcus aureus*. (A-B) Growth inhibition of *S. aureus* (WT). Growth curves of *S. aureus* (WT) treated with respective compounds (200 µM) were recorded at 37°C for 24 h (A). Subsequently, cell viability relative to the wild-type was assessed through spot dilution series (B). - = DMSO control, Van = Vancomycin. In (A), data are represented as mean, n = 3. In (B), data are represented as mean ±SD, n = 3. (C) Growth curves of *S. aureus* (WT) and *S. aureus* knockout strains (*clpC::erm* and *ΔclpP*) treated with respective compounds (200 µM) were recorded at 37°C for 20 h. ns = not significant, \**p* < 0.05, \*\**p* < 0.01, \*\*\**p* < 0.001 by one-way ANOVA with Dunnett’s post-hoc test (B).

Together, our data indicate that although all compounds effectively activate ClpC *in vitro*, they are not yet efficient to cause ClpC-dependent toxicity in *Staphylococcus aureus* and need further optimization.

## Discussion

In this study, we identify and characterize small molecules that deregulate the ClpC/ClpP protease of *Staphylococcus aureus*. Using a high-throughput biochemical screen and thorough validation, we demonstrate that *S. aureus* ClpC can be chemically activated by drug-like small molecules. Eight such compounds significantly enhance ClpC-dependent ATP hydrolysis and proteolysis *in vitro*, triggering adaptor-independent and deregulated protein degradation. Such deregulation of an AAA+ protease is expected to create toxicity *in vivo*, allowing for targeting these proteolytic machines even though they are typically not essential for bacterial viability. All eight compounds target the ClpC N-terminal domain (NTD), underlining its central role in integrating diverse stimuli, including substrates and adaptors, to regulate the AAA+ protein.

Six of the validated compounds bind the hydrophobic substrate-binding groove of the ClpC NTD, a site homologous to the cyclic-peptide binding pocket in mycobacterial ClpC1. A co-crystal structure obtained with Cpd-8 confirms engagement of this groove and reveals specific protein–ligand interactions underlying Cpd-8 mediated ClpC activation. The Cpd-8 binding mode deviates from the ones previously described for the cyclic peptides. However, their similar binding site and functional outcome suggest that the NTD hydrophobic groove represents a conserved regulatory hotspot across ClpC homologs, highlighting this site as a broadly exploitable node for chemical intervention in ClpC activity control. The determined Cpd-8 binding site, together with docking analyses, functional mutagenesis, and structure–activity relationship (SAR) studies based on Cpd-8 derivatives, support a model in which hydrophobic-groove ligands mimic aspects of adaptor (MdfA) (Massoni et al. 2025) or substrate engagement. Cpd-8 putatively destabilizes the inactive ClpC resting state by steric clashes of either the ClpC N-terminal tail with the AAA1 domain or Cpd-8 itself. Further exploration of the co-crystal structure suggests opportunities for structure-guided optimization. In particular, extending Cpd-8a toward the key regulatory residue K85 could enhance both affinity and efficacy.

In addition to the hydrophobic groove, our study identifies the pArg1 pocket as a second, mechanistically distinct site that can be engaged by small molecules to activate ClpC. The pArg1 site recognizes phosphorylated arginines as degrons and functions as allosteric pocket linking degron binding to ClpC activation (Jenne et al. 2025; Trentini et al. 2016). While BacPROTACs exploit this site through pArg-containing ligands (Morreale et al. 2022), engagement of the pArg1 pocket by classical small molecules had not been demonstrated. Here, *in silico* docking predictions, targeted mutagenesis, biochemical assays, and structure–activity relationship analysis of Cpd-10 and its derivatives provide converging evidence that this pocket is druggable and that its occupation by small molecules is sufficient to induce ClpC activation. Presumably, Cpd-10 activates ClpC in a manner similar to how arginine-phosphorylated substrates destabilize the resting state, by clashes with the tip of the regulatory middle domain, as previously suggested (Jenne et al. 2025). These findings extend the functional repertoire of the pArg1 site and establish it as a viable target for chemical modulation.

While all compounds induce ClpC/ClpP proteolytic activity, they do not lead to the formation of canonical protease complexes. Negative-stain electron microscopy revealed the formation of heterogeneous ClpC/ClpP assemblies upon compound binding. The precise architecture of these complexes varies, however, they are composed of common building blocks including ClpC ring dimers and tetrahedrons among others. Higher-order ClpC/ClpP assemblies were observed irrespective of whether compounds targeted the hydrophobic groove or the pArg1 site. Higher-order assemblies composed of multiple ClpC hexamers have been reported to form in the presence of the toxic cyclic peptide cyclomarin A (Taylor et al. 2022; Maurer et al. 2019), pArg-substrates (Morreale et al. 2022), or derepressed ClpC mutants in the presence of the substrate FITC-casein (Jenne et al. 2025), respectively. These observations suggest that different regulatory sites all share a common ClpC activation pathway that is closely linked to assembly formation.

An important feature of the small-molecule activators described here is their capacity to sustain ClpC–ClpP proteolytic activity over time. Unlike the adaptor protein MecA, which is rapidly degraded as part of an intrinsic negative feedback loop on ClpC activation, these compounds are not depleted in the process. Consequently, they enable longer-lasting protease activity and higher overall substrate breakdown. This highlights a fundamental difference between natural adaptor regulation and chemical deregulation, with promising implications for the controlled manipulation of bacterial proteostasis. Notably, *M. tuberculosis* encodes for ClpC2 and ClpC3, which harbor domains homologous to the ClpC1 NTD. ClpC2 and ClpC3 can sequester cyclic peptides (Hoi et al. 2023; Taylor et al. 2023), thereby protecting ClpC1 from drug-induced deregulation. Such a mechanism is not expected to exist for compounds targeting *S. aureus* ClpC.

Despite robust biochemical activation of ClpC, the compounds exhibited antibacterial activity against *S. aureus* that was independent of ClpC or ClpP in cellular assays. While this precludes a direct link between ClpC activation and growth inhibition in *S. aureus*, it does not detract from the mechanistic insights provided by this study. Rather, it highlights the challenges inherent in translating *in vitro* protease modulation into cellular phenotypes and underscores the need to further optimize selectivity and cellular engagement. In the case of the most toxic Cpd-10, the observed ClpC-independent toxicity may reflect off-target effects mediated by its aesculetin moiety, which is a coumarin derivative. Interestingly, aminocoumarins are natural antibiotics that inhibit DNA gyrase (Vanden Broeck et al. 2019). Although these potential off-target interactions were not directly assessed here, they provide a plausible explanation for the observed cellular growth inhibition. Given the dual activating and inhibiting nature of Cpd-10 on ClpC activity and the fact that inhibition of ClpC1 is toxic in mycobacteria, it would be interesting to evaluate Cpd-10 and related derivatives in mycobacterial systems, where engagement of the essential ClpC1 homologue may yield distinct cellular outcomes.

More broadly, our findings contribute to a growing perception of AAA+ proteases as chemically addressable molecular machines whose activity can be tuned through allosteric regulation. By mapping two distinct ligandable sites within the ClpC NTD and linking their engagement to a defined functional outcome, this work advances the understanding of ClpC activity control mechanisms and expands the toolkit for chemical biology approaches to target this bacterial PQC component. These insights may inform future efforts to design chemical compounds or degradation strategies targeting ClpC.

### Limitations of the study

While the small molecules identified in this study effectively activate *S. aureus* ClpC *in vitro*, their *in vivo* effectiveness appears to be ClpC/ClpP-independent. This discrepancy may stem from reduced uptake due to membrane impermeability or off-target effects. Additionally, their relatively low to moderate affinity may not be sufficient to significantly disrupt proteostasis by deregulating ClpC *in vivo*. Clarifying the first issue could involve measuring intracellular compound concentrations, and off-targets could be identified using techniques like photo-crosslinking probes. Furthermore, enhancing binding affinities to ClpC, guided by structural insights into SARs and the proposed MoA for Cpd-8 and Cpd-10, should be a priority to optimize compounds and achieve ClpC-dependent toxicity while minimizing off-target effects. In support of such attempts, further structural information about the respective compound binding modes is required, as most of the evidence provided here is indirect and mostly based on *in silico* prediction models.

Moreover, our study did not investigate whether these compounds reduce biofilm formation and virulence in a host-infection model, which would provide valuable insights into their therapeutic potential. Thus, future studies should aim to assess the efficacy of optimized ClpC activators in relevant infection models to determine their potential as antivirulence agents and to better understand the role of ClpC in *S. aureus* pathogenesis.

Regarding Cpd-10, its binding to an unknown second site that might counteract ClpC activation warrants caution and should be considered in future drug development. Elucidation of the nature of this secondary binding site and its potential impact on ClpC activity will be essential for the rational design of improved compounds.

## Methods/Experimental procedures

### Strains and plasmids

All strains and plasmids used in this study are summarized in Table S2. *Escherichia coli* (*E. coli*) cells were cultivated in LB medium at 30°C with agitation at 120 rpm, supplemented with appropriate antibiotics. For protein overexpression, 2x YT medium (16 g/l tryptone, 10 g/l yeast extract, 5 g/l NaCl, adjusted to pH 7) was used. *E. coli* XL1 Blue was utilized for cloning and retaining of plasmids requiring kanamycin at 50 μg/ml and ampicillin at 100 μg/ml for plasmid propagation.

### Protein purification

*Staphylococcus aureus* (*Sa*) ClpC (WT and derivatives), ClpP, and MecA were purified from *E. coli ΔclpB::kan* cells overexpressing them using pDS56-derived expression vectors (Carroni et al. 2017). ClpC deletion mutants were generated by PCR, and point mutants were constructed by Quikchange one-step site-directed mutagenesis. All mutations were verified by sequencing. ClpC, ClpP, MecA, FtsZ, MecA-mEOS3.2, and isolated ClpC-NTD harbor a C-terminal His6-tag and were purified using Ni-IDA (Macherey-Nagel) following the instructions provided by the manufacturer. In short, cell pellets were resuspended in buffer A (50 mM NaH_2_PO_4_, 300 mM NaCl, 5 mM β-mercaptoethanol, pH 8.0) supplemented with protease inhibitors (Roche). After cell lysis using a French press, the cell debris was removed by centrifugation at 18,000 g for 60 min at 4°C, and the cleared lysate was subsequently transferred into a plastic column containing pre-equilibrated 0.8-1 g Protino™ Ni-IDA resin (Macherey-Nagel) at 4°C. Afterwards, the resin was washed once with buffer A. His-tagged proteins were eluted by adding buffer A supplemented with 250 mM Imidazole. Subsequently, pooled protein fractions were subjected to size exclusion chromatography (Superdex S200, Amersham) in buffer B (50 mM Tris-HCl, pH 7.5, 50 mM KCl, 20 mM MgCl_2_, 5% (v/v) glycerol, 2 mM DTT).

Pyruvate kinase of rabbit muscle and Casein fluorescein isothiocyanate (FITC-casein) from bovine milk were purchased from Sigma. Protein concentrations were determined with the Bradford assay (BioRad).

### High-throughput screen (HTS)

337 assay plates (384-well format) were screened in 7 batches. These plates are pre-plated with compounds in columns 3 to 22 (1 µl of 400 µM compound solution in 4% DMSO) and 1 µl of 4% DMSO solution in columns 1 and 2. In columns 1 to 22, 4 µl of ClpP/ClpC reaction mix was added using a MultiDrop device (ThermoFisher Scientific); in columns 23 and 24, the 4 µl of reaction mix contained MecA as well and served as the positive control. The reaction was started by adding 5 µl of an ATP solution (Buffer: 50 mM Tris pH 7.5, 20 mM KCl, 10 mM MgCl_2_, and 2 mM DTT; ATP, Phosphoenolpyruvate (PEP), and pyruvate kinase (PK)) with a FlexDrop (Revvity, with plate stacker) to a final assay volume of 10 µl. The final ingredient concentrations in each well were as follows: 0.4% DMSO, 0.5 µM ClpC, 0.75 µM ClpP, 0.5 µM FITC-casein, 2 mM ATP, 3 mM PEP, 200 ng PK, and 0.75 µM MecA in the positive control wells. Compounds were screened at a single concentration (40 µM). After 60 minutes of incubation, the fluorescence signal (Excitation 485 nm / Emission 535 nm) was read with the Envision HTS reader from Revvity (with plate stacker). For each batch, 2 QC-plates (plates with no screening compounds in column 3-22) were screened.

### Biochemical assays

Buffer C (50 mM Tris-HCl, pH 7.5, 50 mM KCl, 20 mM MgCl_2_, 2 mM DTT) was used for all biochemical assays. Compounds were added to a final concentration of 50 µM with 1% (v/v) DMSO unless stated otherwise.

### ATPase assay

Rates of ATP hydrolysis were determined using a coupled-colorimetric assay as described previously (Oguchi et al. 2012). Oxidation of NADH to NAD^+^ was monitored by measuring absorbance at 340 nm.

ATPase activity measurements were performed at final concentrations of 0.5 μM ClpC, 0.75 μM ClpP, 0.75 μM MecA, 0.5 μM FITC-cas, and an ATP regenerating system (PK/LDH mix: 250 µM NADH, 500 µM PEP, 1/20 (v/v) PK/LDH (Sigma-Aldrich)) in buffer C supplemented with 2 mM ATP at 30°C in a transparent 96-well plate (Greiner) using a CLARIOstar^plus^ plate reader (BMG Labtech). Isolated ClpC-NTD was added to a final concentration of 25 µM.

The ATP hydrolysis rate for at least three biological replicates was calculated according to the following equation:

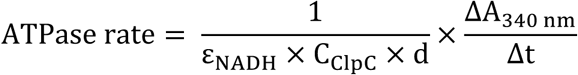

ε_NADH_ – molar absorption coefficient of NADH at a wavelength of 340 nm (6220 M^-1^cm^-1^).

C_ClpC_ – final concentration of ClpC (0.5 μM).

d – optical path length (1 cm).

ΔA_340 nm_/Δt – slope of the linear decline in absorption at a wavelength of 340 nm.

### Degradation assays

Proteolytic activity was determined using fluorescein-labelled casein (FITC-casein) and FtsZ at final concentrations of 0.5 μM ClpC, 0.75 μM ClpP, 0.75 μM MecA, 0.5 μM FITC-casein, 1 µM FtsZ, 0.5 µM MecA-mEOS3.2, and an ATP regenerating system (3 mM PEP, 20 ng/μl PK (Sigma-Aldrich)) in buffer C. Isolated ClpC-NTD was added to a final concentration of 25 µM. ADEP1 was used in FITC-casein degradation assays at a concentration of 10 μg/ml. Reactions were started by the addition of ATP to a final concentration of 2 mM.

The increase of fluorescence upon FITC-casein degradation was monitored at 483-14 nm and 530-30 nm as excitation and emission wavelengths, respectively, in black 96-well plates (Greiner) using a CLARIOstar^plus^ plate reader (BMG Labtech). The increase of fluorescence upon MecA-mEOS degradation was monitored at 489-14 nm and 529.5-17 nm as excitation and emission wavelengths, respectively, in black 96-well plates (Greiner) using a CLARIOstar^plus^ plate reader (BMG Labtech). The raw fluorescence signal was normalized by setting the initial value to 100%. The FITC-casein fluorescence gain (% min^-1^) was calculated by determining the initial slopes of the linear fluorescence signal increase for at least three biological replicates. Cpd-10 presence decreased total FITC-casein fluorescence.

FtsZ degradation was monitored by SDS-PAGE followed by Coomassie staining. Band intensities were quantified using FIJI (Schindelin et al. 2012).

### Analytical size-exclusion chromatography (SEC)

Complex formation of ClpC and respective variants was monitored via analytical size-exclusion chromatography. SEC was performed at 4°C using an ÄKTA pure (Cytiva) equipped with a Superose 6 Increase 10/300 GL column (Cytiva), pre-equilibrated with buffer B supplemented with 2 mM ATP. 6 μM ClpC-E280A/E618A (ClpC-DWB) in the presence of 9 μM ClpP and 6 μM FITC-casein was pre-incubated with 2 mM ATPγS and 50 µM of respective compounds at RT for 5 min, and 200 μl of the sample were loaded onto the column. Aliquots were taken and subjected to SDS-PAGE analysis. Gels were stained using SYPRO Ruby Protein Gel Stain. For each sample containing a compound, two biological replicates were performed.

### Nano differential scanning fluorimetry (nanoDSF)

Domain stability was assessed via nano differential scanning fluorimetry (nanoDSF). 100 μM ClpC-NTD-WT and L14D-I88R in 50 mM HEPES (pH 7.5), 50 mM KCl, and 2 mM DTT were loaded into Prometheus^TM^ high sensitivity capillaries (NanoTemper). Thermal unfolding was analyzed by gradually heating from 20°C to 95°C at a rate of 0.5°C/min with a Prometheus^TM^ Panta device (NanoTemper), while tracking turbidity and the intrinsic fluorescence at 330/350 nm. The melting temperature (T_m_) was determined using the Prometheus Panta Analysis software (version 1.4.3). Three technical replicates were performed for each sample.

### Negative stain electron microscopy (nsEM)

Negative staining, data collection, and processing were performed as described previously (Ohi et al. 2004). In brief, a 5 µl sample was applied to a glow-discharged grid with a 6-8 nm thick layer of continuous carbon. After incubation for 5-60 s, the sample was blotted on a Whatman filter paper 50 (1450-070) and quickly washed with three drops of water. Samples on grids were stained with 2.5 % aqueous uranyl acetate. Images were acquired using a ThermoFisher Talos L120C electron microscope equipped with a Ceta 16M camera, operated at 120 kV. The micrographs were acquired at ×36,000 (Cpd-8, Cpd-10) or ×57,000 (Cpd-41) magnification (resulting in 4.13/2.55 Å per pixel, respectively) using EPU software. For 2D classification of ClpC-DWB + ClpP + FITC-casein + Cpd-8, 6,925 particles were selected using the boxing tool in EMAN2 (box pixel size 360) (Tang et al. 2007). Image processing was carried out using the IMAGIC-4D package (van Heel et al. 1996). Particles were band-pass filtered, normalized for gray value distribution, and mass centered. 2D alignment, classification, and iterative refinement of class averages were performed as previously described (Liu and Wang 2011). Following classification, particles were sub-classified to further improve the data quality of selected classes.

Samples were prepared for and subjected to analytical SEC as described previously, with the peak fraction of the early eluting large complexes being applied to a grid.

### *In vivo* toxicity assays

*S. aureus* strains (WT: ATCC 25923, *clpC::erm*/*ΔclpP*: NCTC 8325-4) were cultured on Columbia Agar plates containing 5% Sheep Blood (BD Biosciences). For assay preparations, single colonies were transferred into trypticase soy bouillon (TSB; BD, Germany) and grown overnight at 37°C with shaking at 150 rpm. The following day, the bacterial cells were diluted to an OD_600_ = 0.05, grown in TSB until reaching exponential growth (OD_600_ = 0.6-0.8), and then diluted to a final concentration of 1 x 10^5^ CFU/ml in a polystyrene U-bottom 96-well plate (Greiner) containing respective compounds (10/50/200 µM, as specified) at a final DMSO concentration of 2% (v/v). The plate was sealed with a gas-permeable Breathe-Easy® sealing membrane (Sigma). Wells supplemented with Vancomycin and DMSO were included as control groups, respectively. Growth curves (OD_600_) were recorded at 37°C with shaking at 600 rpm (linear shaking) for 20-24 h using a Victor Nivo microplate reader (Perkin Elmer). Subsequently, the bacterial cells were adjusted to OD_600_ = 1, and serial dilutions (10^-1^ - 10^-6^) were spotted on Columbia Agar plates containing 5% Sheep Blood using a custom-made spotter (ZMBH workshop, University of Heidelberg) to assess cell viability. Plates were incubated overnight at 37°C. Three biological replicates were performed.

### Crystallization, data collection, model building, and refinement

Crystals were grown at 291.15 K using sitting drop vapor diffusion. The crystallization reservoirs were composed of 2.145 M (NH4)_2_SO4, 0.1 M MES pH 6 (ClpC-NTD apo) and 1.909 M (NH4)_2_SO4, 0.1 M Tris pH 8 (ClpC-NTD:Cpd-8). Crystallization drops contained 100 nL reservoir solution and 500 nL protein at a concentration of 7.5 mg/ml. Crystals grew within 14 to 21 days. Crystals were frozen in liquid nitrogen using glycerol as cryo-protectant. Data were collected at ESRF beamline ID30A-1 at cryogenic conditions, integrated using XDS (Kabsch 2010), and scaled using AIMLESS (Evans and Murshudov 2013) as part of the CCP4 software package (Winn et al. 2011). Phases were obtained by molecular replacement with PHASER (McCoy et al. 2007) implemented in the Phenix package (Liebschner et al. 2019) using an AlphaFold2 (Jumper et al. 2021) prediction as search model. The resulting data sets had space group P4_3_2_1_2, a maximum resolution of 1.78 Å (ClpC-NTD apo) and 1.9 Å (ClpC-NTD:Cpd-8) with one ClpC-NTD molecule per asymmetric unit each. Cpd-8 was parametrized from SMILES description using phenix.elbow (Moriarty et al. 2009). Iterative model building and refinement were performed with Coot (Emsley et al. 2010) and phenix.refine (Afonine et al. 2012). The quality of the resulting structural models was analyzed with MolProbity (Chen et al. 2010). Figure S7A was prepared with PyMol 3.1.6.1 (The PyMol Molecular Graphics System, Schrödinger). Figure S7B was prepared using Ligplot+ v2.3 (Laskowski and Swindells 2011). Crystallographic data are summarized in Table S1. Coordinates and structure factors of the two x-ray structures reported in this study are deposited at the Protein Data Bank PDB under accession codes 9TXC (ClpC-NTD apo) and 9TXD (ClpC-NTD:Cpd-8).

### *In silico* predictions

#### DiffDock-L screening for binding site prediction

In order to locate the binding sites and pre-screen for plausible binding poses, we docked all compounds to the N-terminal domain (NTD) of ClpC (apo-ClpC) using DiffDock-L (Corso et al. 2022, 2024). DiffDock is a diffusion-based generative model with a graph neural network architecture trained to predict docking poses given a protein target and a ligand. Although DiffDock can generate poses that violate physical constraints (Buttenschoen et al. 2024) and is not trained with explicit energy or force-field terms, it remains suited for an initial prescreening stage. Its learned generative and scoring model captures patterns of geometric complementarity between ligands and protein pockets, which can be used for the identification of structurally plausible binding regions.

For each compound, we sampled 40 binding poses with the locally installed DiffDock-L (github commit: 9a22cbc) using 40 diffusion timesteps. Most of DiffDock-L predictions converged to three known binding sites in the ClpC-NTD: pArg1, NTD, and pArg2. We then quantified how often DiffDock-L positioned ligands in each of the three sites. For every sampled pose, we computed the center of mass (CoM) of a ligand and computed its Euclidean (L2) distance to the site-defining residue (E106, I88/M92, E32) sidechain atoms. Each pose was assigned to the site whose sidechain atoms were closest (minimum heavy-atom distance); poses with a minimum distance > 10 Å to all side-chain atoms were excluded from further analyses.

#### Biased flexible docking of ligands with HADDOCK 2.4

To obtain physically consistent ligand binding modes, we performed restrained docking with HADDOCK 2.4 using experimental restraints and binding sites obtained with DiffDock-L. For each ligand, we specified a set of active residues (pArg1: T7/E106, Hydrophobic groove: Y80/M92, pArg2: E32) to bias sampling towards the intended binding site. Docking was run in three stages: 1000 structures for rigid-body docking, 200 structures for semi-flexible refinement, and 200 structures for final water refinement. All parameters were kept at their default values, as defined by the developers of HADDOCK, except the electrostatics weight in the HADDOCK score computation, which was set to 0.1. Each ligand was independently docked to all three candidate binding sites, and the median HADDOCK score was computed for the resulting clusters. Final poses shown in the main text were selected based on the total HADDOCK score and visual inspection.

#### Bioinformatic analyses

Structural figures were prepared with ChimeraX v1.10 (Goddard et al. 2018; Pettersen et al. 2021). Chemical structures were prepared using ChemDraw v22.2.0. Curve fitting and statistical analyses were performed using GraphPad Prism v10.6.1 (GraphPad Software). Because the positive control was used as a reference for assay validation, statistical comparisons involving it were conducted separately from the comparisons of compound treatment versus the negative control using unpaired Students’ t-tests.

## Data availability

Coordinates and structure factors of the two x-ray structures reported in this study are deposited at the Protein Data Bank PDB under accession codes 9TXC (ClpC-NTD apo) and 9TXD (ClpC-NTD:Cpd-8).

## Acknowledgments

T.J. was supported by the Heidelberg Biosciences International Graduate School (HBIGS). This work was supported by a grant of the Deutsche Forschungsgemeinschaft to A.M. (MO970/8-1). We acknowledge the support by the grant “Compound Screening” (ExU 6.1.5) by the Flagship Initiative “Engineering Molecular Biosystems” of Heidelberg University. I.S., M.S., and J.K. acknowledge the service SDS@hd, supported by the Ministry of Science, Research and Arts Baden-Württemberg, and the DFG through grant INST 35/1503-1 FUGG. We acknowledge access to ESRF Grenoble beamline ID30A-1 (MASSIF-1) and thank the ESRF staff for their support. We thank Britta Klem from the CCTP/BZH crystallization platform for excellent technical support. We are grateful for access to the infrastructure and support provided by the Cryo-EM Network at Heidelberg University (HDcryoNet). We thank Dorte Frees for providing the *S. aureus* knockout strains used in this study. Lastly, we thank Alexander Dalpke for generously providing laboratory space and infrastructure to conduct S2 experiments.

## Author contributions

Conceptualization, T.J., B.B., and A.M.

Methodology, Investigation, T.J., V.V., B.J., M.S., J.K., D.F., and A.M.

Formal Analysis, T.J., V.V., U.U., M.S., J.K., D.F., and A.M.

Resources, U.U., J.K., D.F., E.S., I.S., B.B., and A.M.

Writing – Original Draft, T.J.

Writing – Review and Editing, T.J., V.V., U.U., J.K., D.F., E.S., and A.M.

Supervision, A.M.

Visualization, T.J., V.V., J.K., and A.M.

Funding Acquisition, B.B. and A.M.

