## Supplemental information for "Small-molecule activators of the *Staphylococcus aureus* ClpC/ClpP AAA+ protease" (from another version of this preprint: version 1, 2026-04-09)

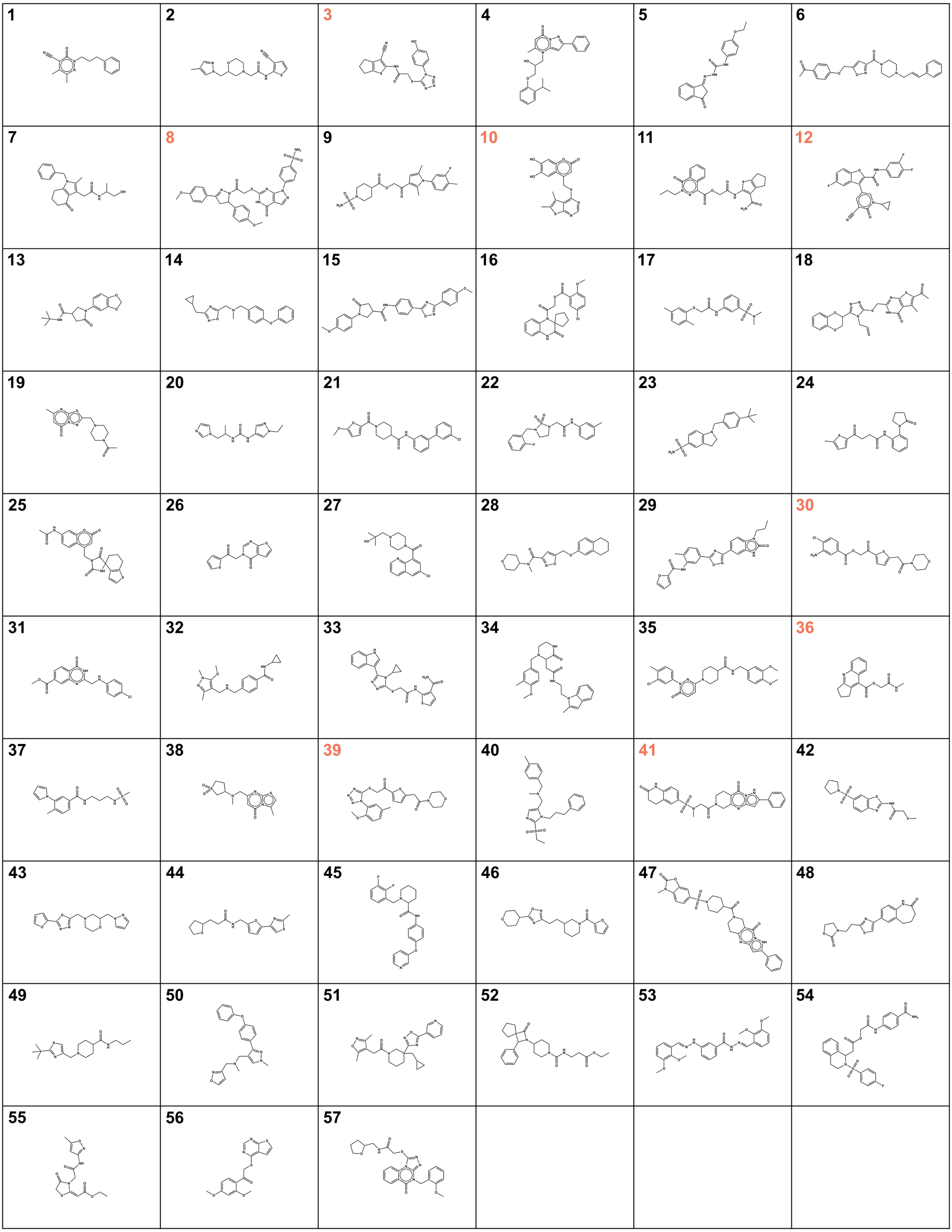

**Figure S1**

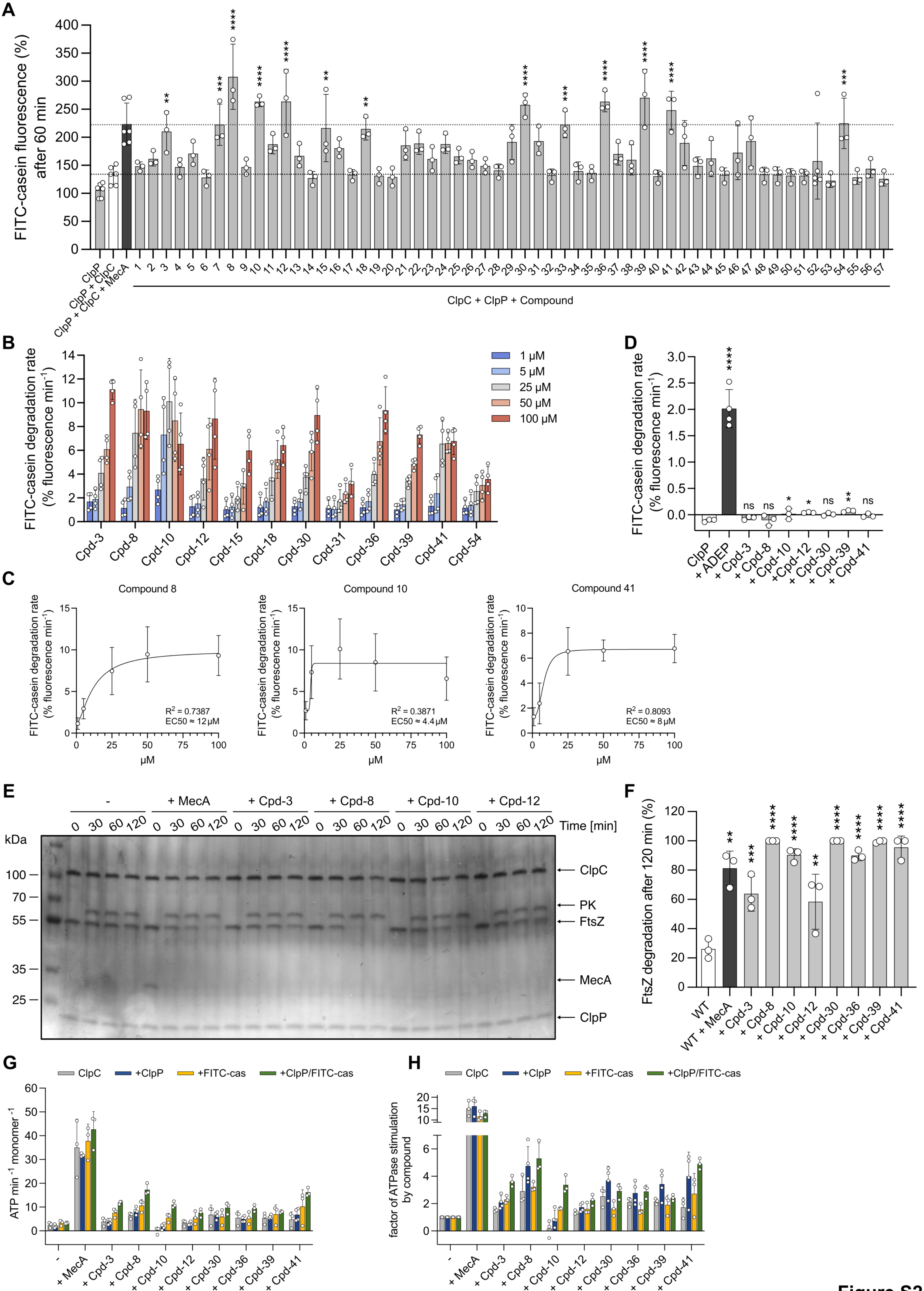

A
FITC-casein fluorescence (%) after 60 min
ClpP
ClpP + ClpC
ClpP + ClpC + MecA
ClpC + ClpP + Compound
B
FITC-casein degradation rate (% fluorescence min-1)
1 µM
5 µM
25 µM
50 µM
100 µM
Cpd-3 Cpd-8 Cpd-10 Cpd-12 Cpd-15 Cpd-18 Cpd-30 Cpd-31 Cpd-36 Cpd-39 Cpd-41 Cpd-54
C
Compound 8
R2 = 0.7387
EC50 ≈ 12 µM
Compound 10
R2 = 0.3871
EC50 ≈ 4.4 µM
Compound 41
R2 = 0.8093
EC50 ≈ 8 µM
µM
D
ClpP
+ ADEP
+ Cpd-3
+ Cpd-8
+ Cpd-10
+Cpd-12
+ Cpd-30
+ Cpd-39
+ Cpd-41
E
- + MecA + Cpd-3 + Cpd-8 + Cpd-10 + Cpd-12
Time [min]
kDa
ClpC
PK
FtsZ
MecA
ClpP
F
FtsZ degradation after 120 min (%)
WT
WT + MecA
G
ClpC +ClpP +FITC-cas +ClpP/FITC-cas
ATP min -1 monomer -1
H
factor of ATPase stimulation by compound

**Figure S2**

A Cpd-8: 2D classes

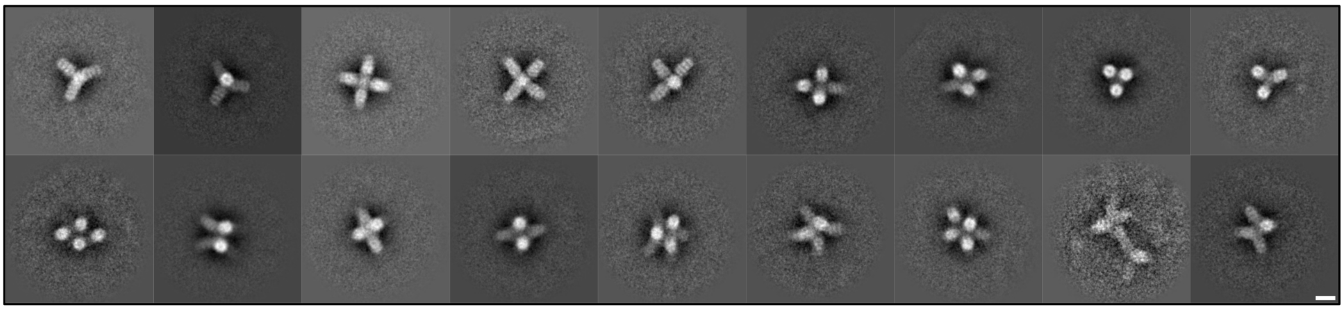

B Cpd-10: singe particles

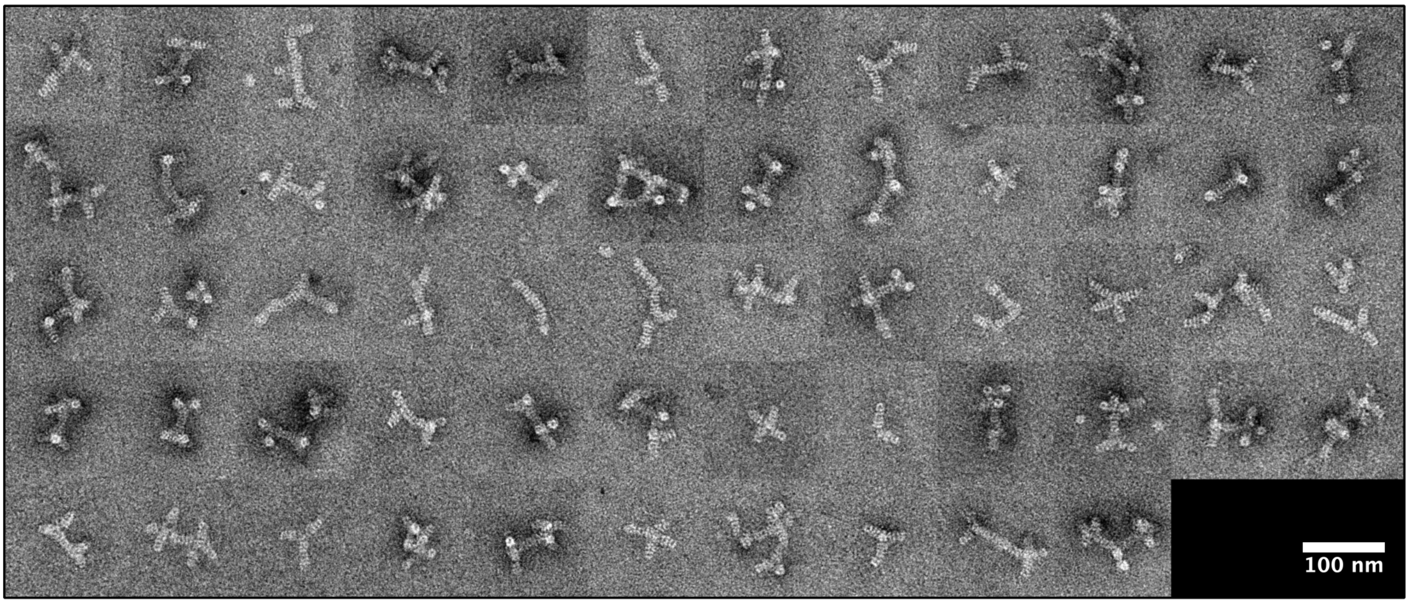

C Cpd-41: single particles

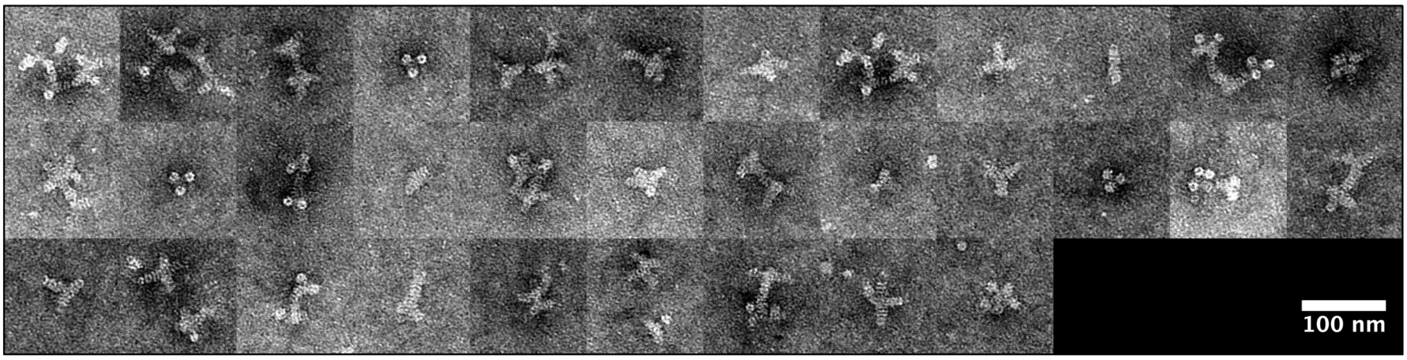

Figure S3

A

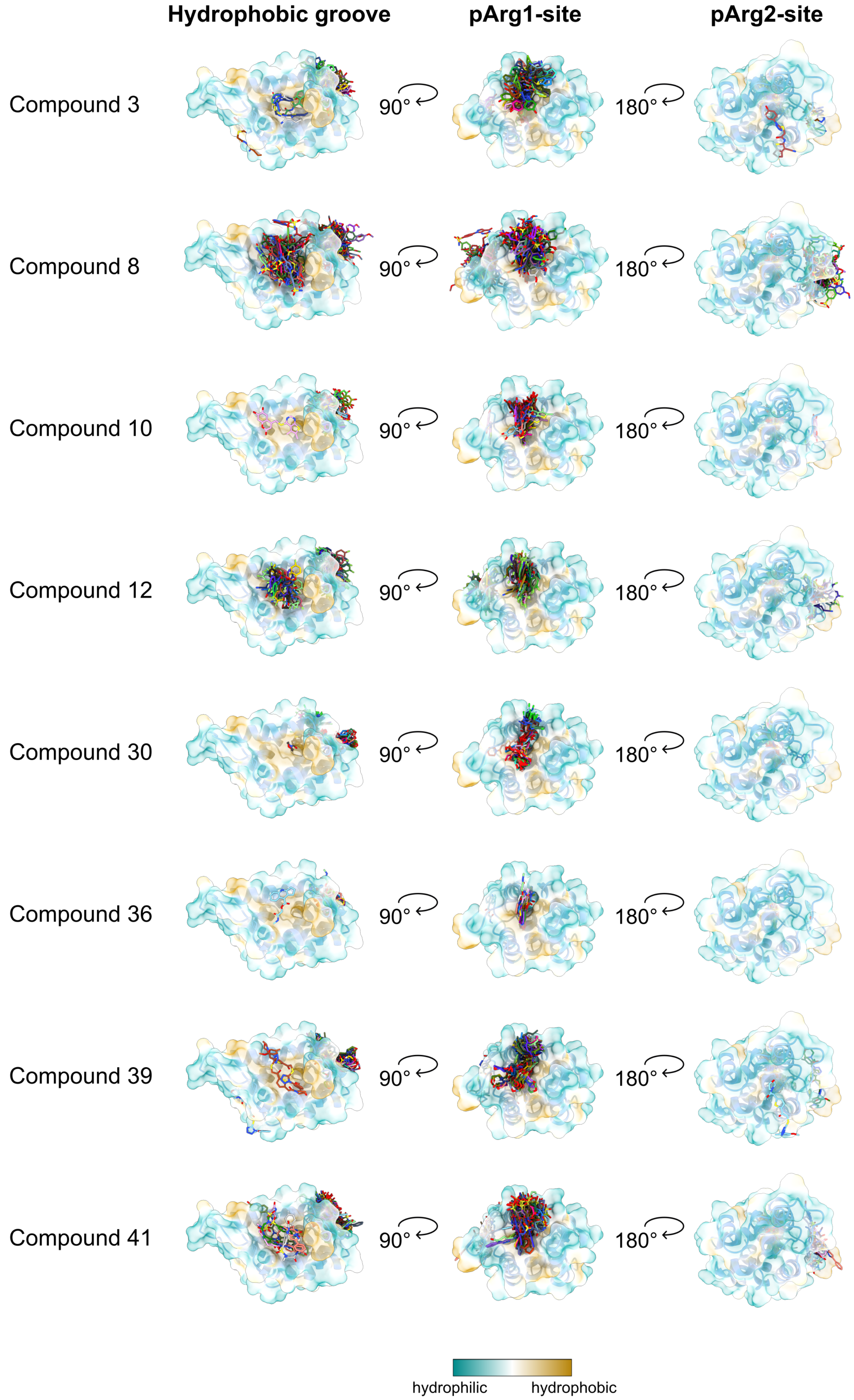

Hydrophobic groove
pArg1-site
pArg2-site
Compound 3
Compound 8
Compound 10
Compound 12
Compound 30
Compound 36
Compound 39
Compound 41
90°
180°
hydrophilic
hydrophobic

**Figure S4**

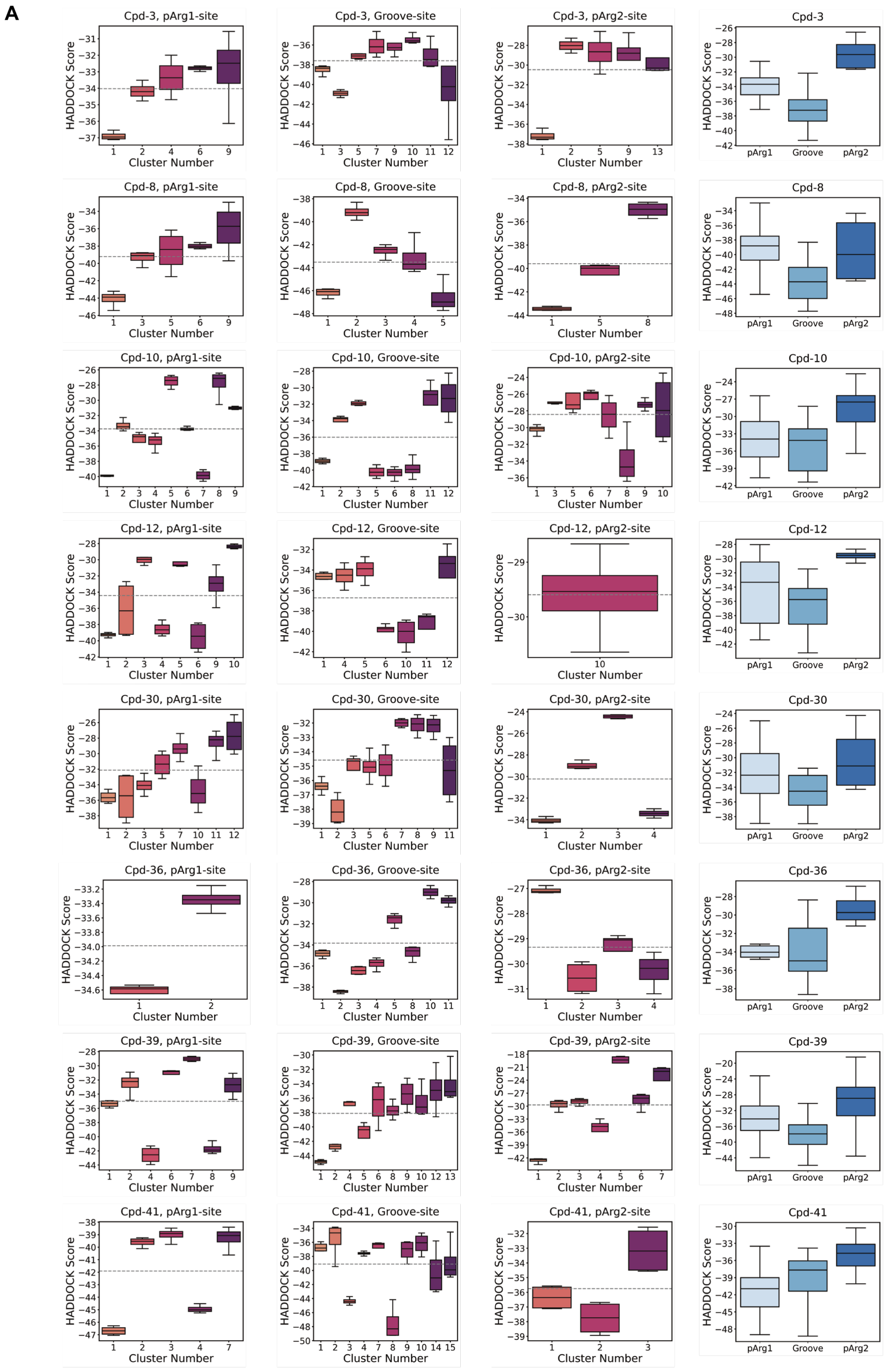
A
Cpd-3, pArg1-site
Cpd-3, Groove-site
Cpd-3, pArg2-site
Cpd-3
Cpd-8, pArg1-site
Cpd-8, Groove-site
Cpd-8, pArg2-site
Cpd-8
Cpd-10, pArg1-site
Cpd-10, Groove-site
Cpd-10, pArg2-site
Cpd-10
Cpd-12, pArg1-site
Cpd-12, Groove-site
Cpd-12, pArg2-site
Cpd-12
Cpd-30, pArg1-site
Cpd-30, Groove-site
Cpd-30, pArg2-site
Cpd-30
Cpd-36, pArg1-site
Cpd-36, Groove-site
Cpd-36, pArg2-site
Cpd-36
Cpd-39, pArg1-site
Cpd-39, Groove-site
Cpd-39, pArg2-site
Cpd-39
Cpd-41, pArg1-site
Cpd-41, Groove-site
Cpd-41, pArg2-site
Cpd-41
HADDOCK Score
Cluster Number
pArg1
Groove
pArg2

Figure S5

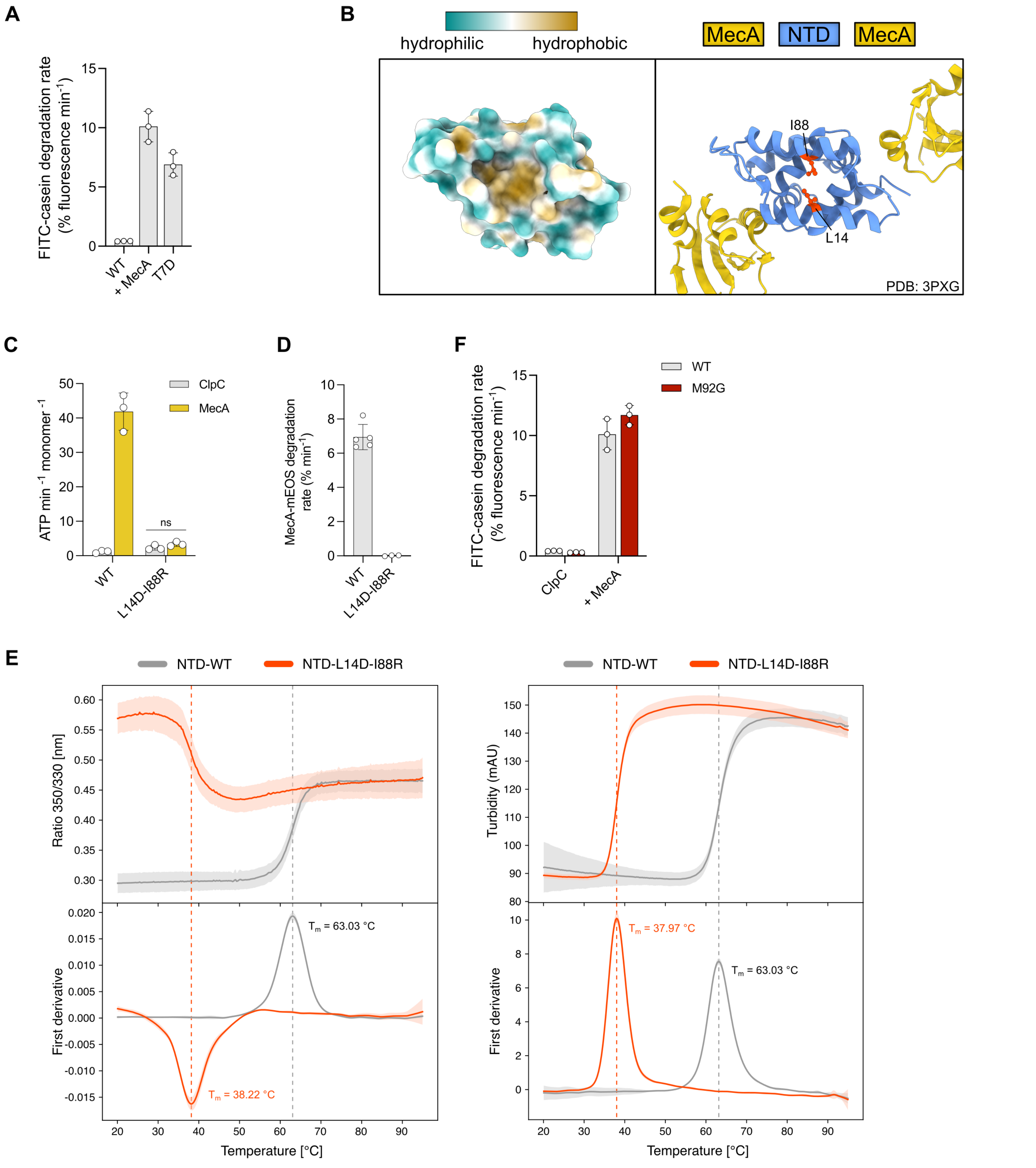

A
FITC-casein degradation rate (% fluorescence min-1)
15
10
5
0
WT
+ MecA
T7D
B
hydrophilic
hydrophobic
MecA
NTD
MecA
I88
L14
PDB: 3PXG
C
ATP min -1 monomer -1
50
40
30
20
10
0
ClpC
MecA
ns
WT
L14D-I88R
D
MecA-mEOS degradation rate (% min-1)
10
8
6
4
2
0
WT
L14D-I88R
F
FITC-casein degradation rate (% fluorescence min-1)
15
10
5
0
WT
M92G
ClpC
+ MecA
E
NTD-WT
NTD-L14D-I88R
Ratio 350/330 [nm]
0.60
0.55
0.50
0.45
0.40
0.35
0.30
First derivative
0.020
0.015
0.010
0.005
0.000
-0.005
-0.010
-0.015
Tm = 63.03 °C
Tm = 38.22 °C
20
30
40
50
60
70
80
90
Temperature [°C]
NTD-WT
NTD-L14D-I88R
Turbidity (mAU)
150
140
130
120
110
100
90
80
First derivative
10
8
6
4
2
0
Tm = 37.97 °C
Tm = 63.03 °C
20
30
40
50
60
70
80
90
Temperature [°C]

**Figure S6**

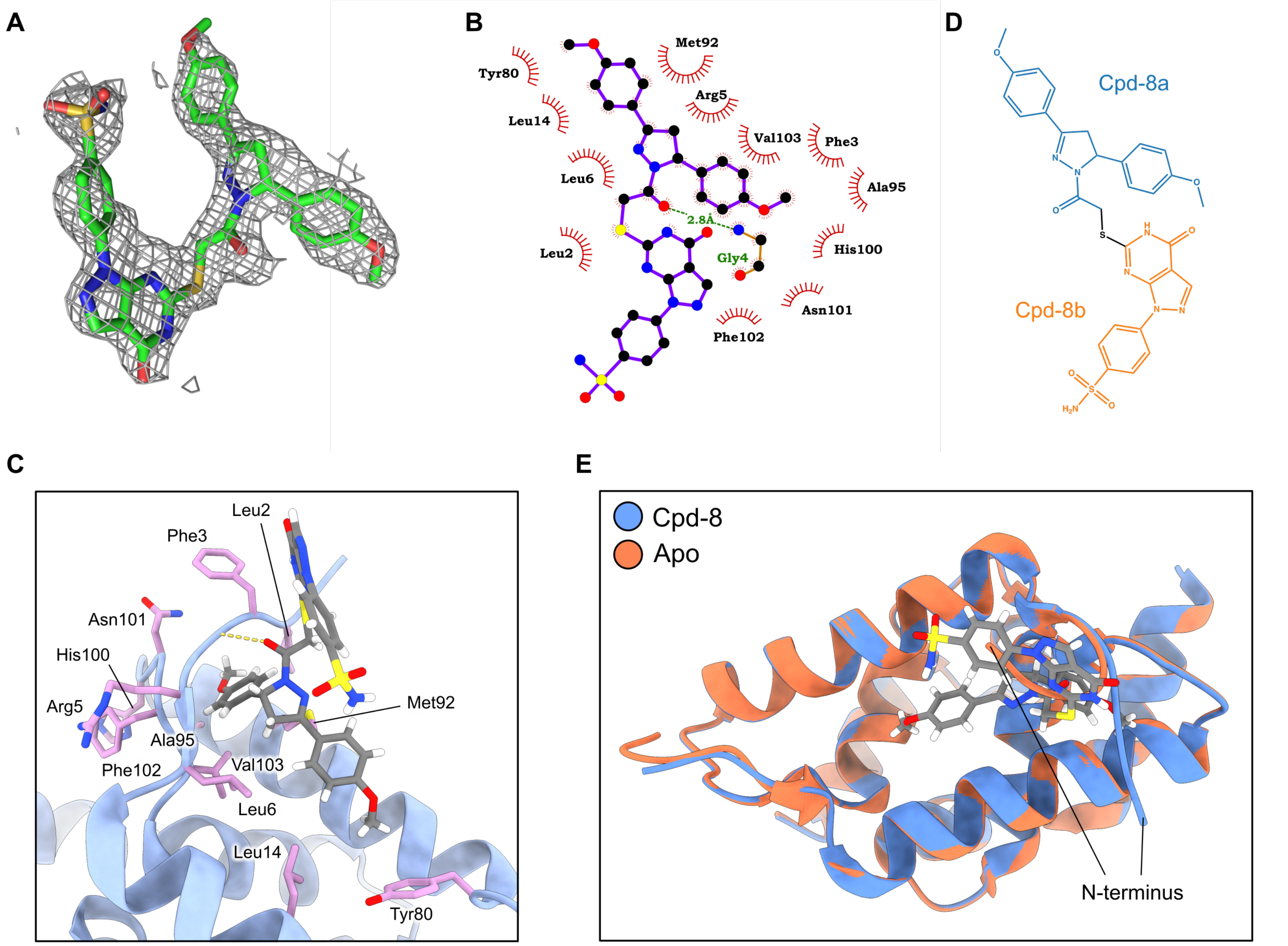
A
B
Tyr80
Met92
Arg5
Leu14
Val103
Phe3
Leu6
Ala95
2.8Å
Leu2
Gly4
His100
Asn101
Phe102
D
Cpd-8a
Cpd-8b
C
Leu2
Phe3
Asn101
His100
Arg5
Ala95
Met92
Phe102
Val103
Leu6
Leu14
Tyr80
E
Cpd-8
Apo
N-terminus

Figure S7

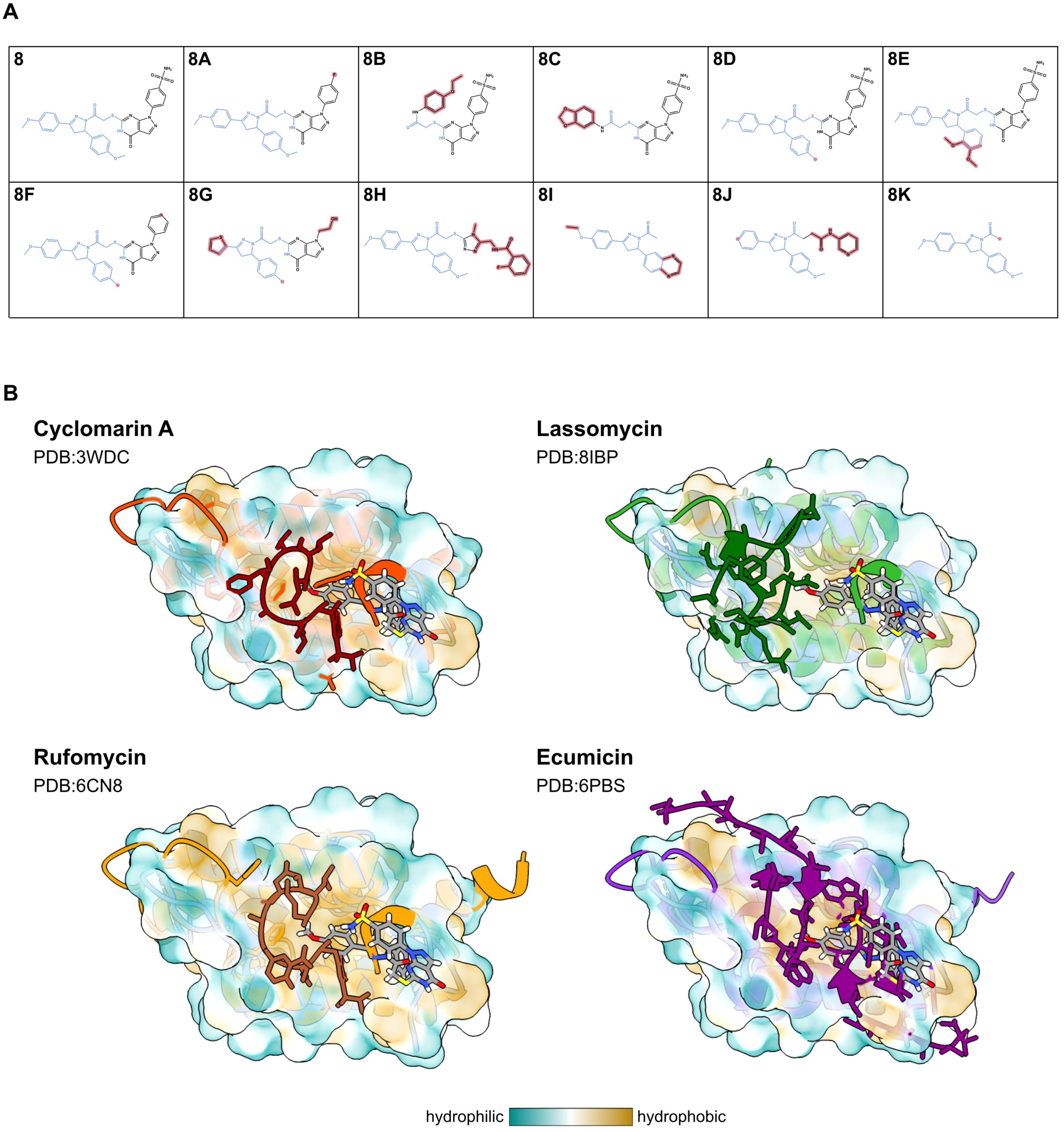
A
8
8A
8B
8C
8D
8E
8F
8G
8H
8I
8J
8K
B
Cyclomarin A
PDB:3WDC
Lassomycin
PDB:8IBP
Rufomycin
PDB:6CN8
Ecumicin
PDB:6PBS
hydrophilic
hydrophobic

Figure S8

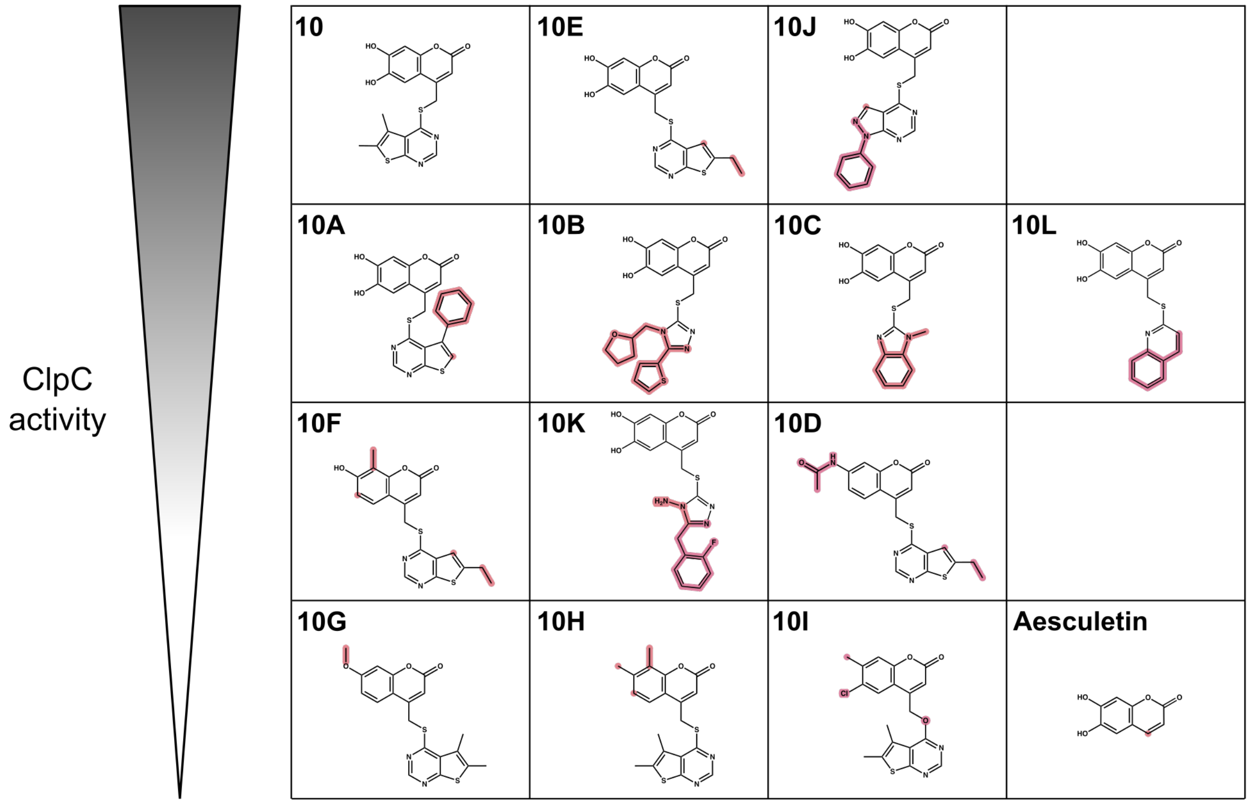
ClpC activity
10
10E
10J
10A
10B
10C
10L
10F
10K
10D
10G
10H
10I
Aesculetin

**Figure S9**

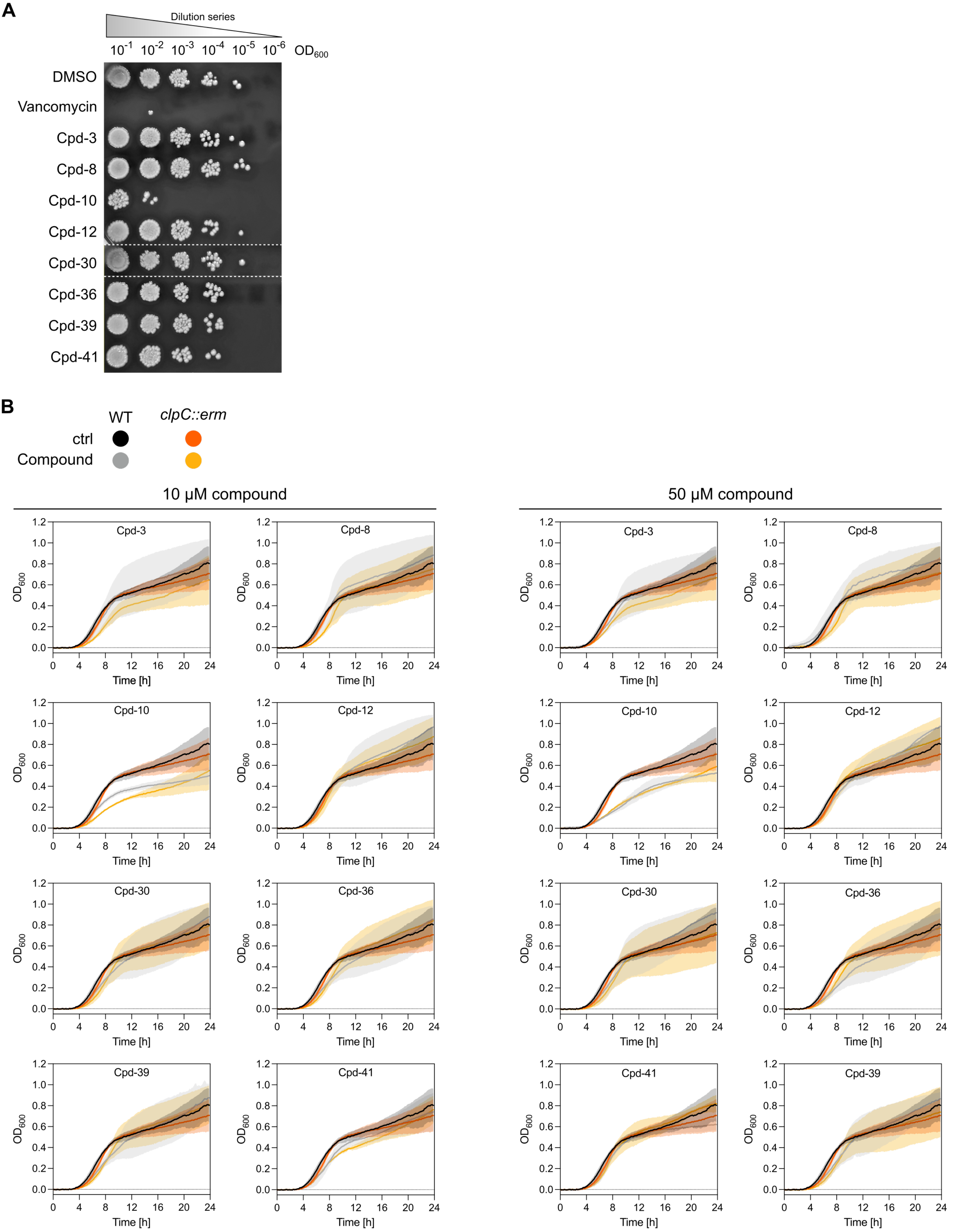
A
Dilution series
10-1 10-2 10-3 10-4 10-5 10-6 OD600
DMSO
Vancomycin
Cpd-3
Cpd-8
Cpd-10
Cpd-12
Cpd-30
Cpd-36
Cpd-39
Cpd-41
B
WT
clpC::erm
ctrl
Compound
10 µM compound
50 µM compound
Cpd-3
Cpd-8
Cpd-10
Cpd-12
Cpd-30
Cpd-36
Cpd-39
Cpd-41
OD600
Time [h]

Figure S10

### Figure legends

#### Figure S1

Chemical structures of identified primary hits. Numbers of compounds selected for extended characterization are highlighted in red.

#### Figure S2

Identified and selected compounds specifically activate ClpC. (A) Total FITC-casein degradation activities after 60 min were determined in the presence of ClpP and ClpC + ClpP ± MecA or respective compounds (50 µM) Initial FITC-casein fluorescence was set to 100%. Data are represented as mean ±SD, n ≥ 3. (B-C) Dose-response curves of ClpC-WT were determined by monitoring FITC-casein degradation activities at increasing compound concentrations in the presence of ClpP (B). Afterwards, EC50 values were obtained by fitting agonist–response data with a four-parameter variable-slope nonlinear regression model (C). Data are represented as mean ±SD, n = 4. (D) FITC-casein degradation activities of ClpP were determined ± ADEP as positive control or respective compounds (50 µM). Data are represented as mean ±SD, n = 3. (E-F) FtsZ degradation activities were determined ± MecA or respective compounds (50 µM) in the presence of ClpP. A representative gel (E) for the quantification of the FtsZ degradation rates (see Figure 1D) and total FtsZ degradation after 120 min (F) is shown. Protein identities are indicated. PK = pyruvate kinase. (G-H) ATPase activities (G) and stimulation factors (H) of ClpC were determined under various conditions (alone, + ClpP, + FITC-casein, and ClpP + FITC-casein) ± MecA or respective compounds (50 µM). Data are represented as mean ±SD, n = 3.

ns = not significant, $^{**}p < 0.01$, $^{***}p < 0.001$, $^{****}p < 0.0001$ by one-way ANOVA with Dunnett's post-hoc test (D, F). Positive controls were assessed separately.

#### Figure S3

Compounds alter ClpC assembly states. (A) Additional 2D classes of Cpd-8 induced ClpC/ClpP assemblies as described in Figure 2C. Scale bar = 20 nm. (B-C) Representative single particles of ClpC/ClpP assemblies induced by Cpd-10 (B) and Cpd-41 (C) revealing high structural heterogeneity. Samples were subjected to EM following SEC as described in Figure 2C.

**Figure S4**

DiffDock simulations predict compounds to bind to the hydrophobic groove or to pArg1-/pArg2-sites of the NTD. DiffDock docking simulations (n = 40) of respective compounds to the ClpC N-terminal domain. The NTD is shown as a surface colored by hydrophobicity, with three orthogonal views highlighting the hydrophobic groove (left), the pArg1 site (middle), and the pArg2-site (right), with all predicted poses displayed for selected compounds.

**Figure S5**

Total HADDOCK score for the corresponding compound docked to each of the indicated binding sites (pArg1, Hydrophobic groove, and pArg2). The grey dashed line indicates a mean across all clusters; the solid line in the boxes indicates the median, and the box outlines the 25th and 75th percentiles, respectively.

**Figure S6**

Mutation of the NTD hydrophobic groove can have destabilizing effects. (A) FITC-casein degradation activities of ClpC-WT ± MecA and ClpC-T7D in the presence of ClpP. Data are represented as mean ±SD, n = 3. (B) Structural representation of ClpC in complex with MecA (Wang et al. 2011), highlighting hydrophobicity of the ClpC-NTD (left) and residues subjected to mutagenesis located at the hydrophobic groove (L14, I88). (C) ATPase activities of ClpC-WT and ClpC-L14D-I88R were determined in presence of ClpP ± MecA. Data are represented as mean ±SD, n = 3. (D) MecA-mEOS degradation activities of ClpC-WT and ClpC-L14D-I88R were determined. Data are represented as mean ±SD, n ≥ 3. (E) The melting temperatures ($T_m$) of NTD-WT and NTD-L14D-I88R were determined via nanoDSF. First derivatives of the 350/330 nm ratio and sample turbidity were determined to calculate $T_m$. Data are represented as mean ±SD, n = 3. (F) FITC-casein degradation activities of ClpC-WT and ClpC-M92G ± MecA in the presence of ClpP. Data are represented as mean ±SD, n = 3. ns = not significant by unpaired t test (B).

**Figure S7**

Crystal structure of the ClpC-NTD with bound Cpd-8. (A) Mesh representation (grey) of the $2mF_{obs}$-$DF_{calc}$ electron density map around the bound ligand Cpd-8 (carbon atoms in green) at a contour level of 1 $\sigma$. (B) Schematic two-dimensional depiction of the interactions between

ClpC-NTD and Cpd-8 created with LigPlot+ (Laskowski and Swindells 2011). The hydrogen bond between Gly4 and Cpd-8 is shown in green, and the residues forming hydrophobic contacts (cutoff ≤ 4Å) are shown as red partial circles. (C) Close-up view of Cpd-8 (carbons in grey) bound to ClpC-NTD (blue) with contacting residues (plum) and the hydrogen bond (yellow dashed line) between Gly4 and Cpd-8. (D) Cpd-8 can be subdivided in two parts: Cpd-8a (blue) and Cpd-8b (orange). (E) Superposition of ClpC-NTD apo (orange) with ClpC-NTD:Cpd-8 (blue/grey) shows the rearrangement of the N-terminus upon binding of Cpd-8.

**Figure S8**

(A) Chemical structures of Cpd-8 and selected derivatives. Structural elements of Cpd-8 involved in protein–ligand contacts are displayed in blue. Modifications of respective derivatives are highlighted in red. (B) The Sa ClpC-NTD:Cpd-8 co-crystal structure was structurally aligned with available co-crystal structures of the Mtb ClpC1-NTD in complex with various cyclic peptides.

**Figure S9**

Chemical structures of Cpd-10 and derivatives. Derivatives are roughly sorted by their stimulatory effects on ClpC/ClpP FITC-casein degradation activities (see Figure 6D). Modifications of respective derivatives are highlighted in red.

**Figure S10**

Compounds induce ClpC/ClpP independent toxicity in *Staphylococcus aureus*. (A) Representative agar plate used to assess cell viability after compound treatment (see Figures 7A and 7B). (B) Growth curves of *S. aureus* WT and *clpC::erm* treated with respective compounds (10 & 50 µM) were recorded at 37°C for 24 h. Data are represented as mean ±SD, n = 3.

***Table S1: Data collection and refinement statistics for the N-terminal domain of S.aureus ClpC-NTD apo and with bound ligand cpd8.*** *Data for highest-resolution shell are in parentheses.*

| | **ClpC-NTD apo** | **ClpC-NTD:Cpd8** |
|---|---|---|
| **Data collection** | | |
| Beamline, wavelength (Å) | ESRF ID30-A1, 0.9677 | ESRF ID30-A1, 0.9677 |
| Space group | $P4_32_12$ | $P4_32_12$ |
| Resolution range (Å) | 38.47-1.78 (1.82-1.78) | 39.51-1.90 (1.94-1.90) |
| Total reflections | 435570 (25508) | 248442 (16388) |
| Unique reflections | 18608 (1031) | 15514 (956) |
| a, b, c (Å) | 60.86, 60.86, 99.29 | 60.75, 60.75, 100.63 |
| $\alpha$, $\beta$, $\gamma$ (°) | 90, 90, 90 | 90, 90, 90 |
| $R_{merge}$ | 0.098 (2.938) | 0.105 (2.852) |
| $R_{pim}$ | 0.021 (0.611) | 0.028 (0.726) |
| Mean ($I/\sigma(I)$) | 18.2 (1.2) | 14.9 (1.2) |
| Multiplicity | 23.4 (24.1) | 16.0 (17.1) |
| Completeness (%) | 100 (100) | 100 (100) |
| $CC_{1/2}$ | 0.999 (0.617) | 0.999 (0.472) |
| **Refinement** | | |
| $R_{work}$ | 0.1911 (0.3716) | 0.1946 (0.2723) |
| $R_{free}$ | 0.2207 (0.3763) | 0.2141 (0.2853) |
| RMSD bond length (Å) | 0.006 | 0.006 |
| RMSD bond angle (°) | 0.700 | 0.730 |
| Ramachandran favored (%) | 98.51 | 99.24 |
| Ramachandran allowed (%) | 1.49 | 0.76 |
| Ramachandran outliers (%) | 0 | 0 |
| Rotamer outliers (%) | 0 | 0 |
| Clashscore | 0.88 | 0.85 |
| Average B-factor (Å$^2$) | 37.6 | 41.2 |
| Protein | 37.2 | 40.4 |
| Ligands | 47.3 | 53.7 |
| Solvent | 40.5 | 39.8 |
| **PDB identifier** | 9TXC | 9TXD |

Table S2: Strains and plasmids used in this study

| Strain | Description | Source or reference |
|---|---|---|
| *E. coli* XL1 blue | *recA1 endA1 gyrA96 thi-1 hsdR1 supE44 relA1 lac* [F' *proAB lacI$^q$ ΔM15* Tn10 (Tcr)] | Stratagene |
| *E. coli* BL21 | *F- ompT lon hsdSB gal dcm λ* (DE3) | Novagen |
| *E. coli ΔclpB* | MC4100 Δ*clpB* (Kanamycin sensitive) | (1) |
| **Plasmid** | **Description** | **Source or reference** |
| pDS56-*clpC* | Vector for IPTG-inducible expression of ClpC-WT in *E. coli ΔclpB* cells | (2) |
| pDS56-*clpP* | Vector for IPTG-inducible expression of ClpP in *E. coli ΔclpB* cells | (2) |
| pDS56-*mecA* | Vector for IPTG-inducible expression of MecA in *E. coli ΔclpB* cells | (2) |
| pDS56-*clpC*-ΔN | Vector for IPTG-inducible expression of ClpC-ΔN (2-146) in *E. coli ΔclpB* cells | (2) |
| pDS56-*clpC*-ΔN-F436A | Vector for IPTG-inducible expression of ClpC-ΔN-F436A in *E. coli ΔclpB* cells | (2) |
| pDS56-*clpC*-E280A/E618A (DWB) | Vector for IPTG-inducible expression of ClpC-DWB (E280A/E618A) in *E. coli ΔclpB* cells | (2) |
| pDS56-*clpC*-T7D | Vector for IPTG-inducible expression of ClpC-T7D in *E. coli ΔclpB* cells | (1) |
| pDS56-*clpC*-M92G | Vector for IPTG-inducible expression of ClpC-M92G in *E. coli ΔclpB* cells | This study |
| pDS56-*clpC*-L14D-I88R | Vector for IPTG-inducible expression of ClpC-L14D-I88R in *E. coli ΔclpB* cells | This study |
| pET24a-*clpC*-NTD (1-155) | Vector for IPTG-inducible expression of ClpC-NTD-WT in *E. coli* BL21 cells | This study |
| pET24a-*clpC*-NTD-L14D-I88R (1-155) | Vector for IPTG-inducible expression of ClpC-NTD-L14D-I88R in *E. coli* BL21 cells | This study |
| pET24a-ftsZ | Vector for IPTG-inducible expression of FtsZ in *E. coli* BL21 cells | (3) |
| pDS56-*mecA-mEOS3.2* | Vector for IPTG-inducible expression of MecA-mEOS3.2 in *E. coli ΔclpB* cells | (4) |
